# Scaling of cellular proteome with ploidy

**DOI:** 10.1101/2021.05.06.442919

**Authors:** Galal Yahya, Paul Menges, Devi Anggraini Ngandiri, Daniel Schulz, Andreas Wallek, Nils Kulak, Matthias Mann, Patrick Cramer, Van Savage, Markus Raeschle, Zuzana Storchova

**Affiliations:** Dept. of Molecular Genetics, TU Kaiserslautern, Paul-Ehrlich-Strasse 24, 67663 Kaiserslautern, Germany; Department of Microbiology and Immunology, School of Pharmacy, Zagazig University, Egypt; Institute of Molecular Biology, University of Zurich, Switzerland; Max Planck Institute of Biochemistry, 82152 Martinsried, Germany; Max Planck Institue of Biophysical Chemistry, Goettingen, Germany; Department of Biomathematics, University of California at Los Angeles, Los Angeles, CA 90095, United States

## Abstract

Ploidy changes are frequent in nature and contribute to evolution, functional specialization and tumorigenesis (1,2). Analysis of model organisms of different ploidies revealed that increased ploidy leads to an increase in cell and nuclear volume, reduced proliferation (2-4), metabolic changes (5), lower fitness (6,7), and increased genomic instability (8,9), but the underlying mechanisms remain poorly understood. To investigate how the gene expression changes with cellular ploidy, we analyzed isogenic series of budding yeasts from 1N to 4N. We show that mRNA and protein abundance scales allometrically with ploidy, with tetraploid cells showing only threefold increase in proteins compared to haploids. This ploidy-specific scaling occurs via decreased rRNA and ribosomal protein abundance and reduced translation. We demonstrate that the Tor1 activity is reduced with increasing ploidy, which leads to rRNA gene repression via a novel Tor1-Sch9-Tup1 signaling pathway. mTORC1 and S6K activity are also reduced in human tetraploid cells and the concomitant increase of the Tup1 homolog Tle1 downregulates the rDNA transcription. Our results revealed a novel conserved mTORC1-S6K-Tup1/Tle1 pathway that ensures proteome remodeling in response to increased ploidy.

---

The majority of eukaryotic organisms are diploid (2N, with two sets of chromosome) or haploid (1N), but polyploid cells (>2N) are common in nature. Polyploidy is found throughout the eukaryotic kingdom and plays an important role in speciation, especially in plants. Here, polyploidy increases adaptive potential, bringing short-term success but also disadvantages, reflected reflected in the fact that the number of established whole-genome duplications (WGDs) is low (10,11). Additionally, polyploidy plays an important role in differentiation of multicellular organisms, where it arises from developmentally tightly controlled polyploidization or in response to stress conditions in specialized organs and tissues (5). Polyploidy can also occur from an error. Unscheduled polyploidy is common in human cancers, and an estimated 37% of all cancers underwent a WGD event at some point during their progression (12); the incidence of WGD is even higher in metastasis (13). In addition, tetraploidy promotes tumorigenesis in cancer model systems (14). WGD generally impairs genome stability (9,15–18), but also increases cell evolvability and adaptability (19) and it is therefore considered an important driving force in evolution and tumorigenesis (1,20). Despite the frequent occurrence of polyploidy and its importance in evolution and pathology, the effects of WGD on cellular physiology are only partially understood.

Budding yeasts *Saccharomyces cerevisiae* of different ploidy can be constructed and serve as an excellent model to study the consequences of WGD. An apparent result of increased ploidy is the linear increase in cell and nuclear volume, accompanied by a nonlinear (1.58-fold) scaling of two-dimensional structures such as membranes, and a 1.26-fold increase in linear structures (2,9). Polyploid budding yeasts grow at slightly reduced rates compared with diploids (2–4), reaching a larger volume in a similar time span. Increased ploidy in yeast leads to aberrant cell cycle regulation and response to nutrition (2,21), lower fitness (6,7), impaired genome stability (8,9) and prominent evolvability (19). The inherent genome instability of tetraploid yeasts leads to a ploidy reduction to diploidy during in vitro evolution, likely due to the loss of individual chromosomes (3,19). Analyses of the available transcriptomes show that the expression of only a handful of genes changes disproportionately in response to ploidy, suggesting that polyploidization largely tends to maintain the balance of gene products (2,9,22). The few genes with altered mRNA abundance encode membrane and cell wall proteins, likely reflecting an adaptation to the lower surface-to-volume ratio in larger polyploid cells (2,22). The minimal changes observed in transcriptome raised the question of whether ploidy-dependent regulation occurs post-transcriptionally. Therefore, we set out to analyze proteome changes in response to altered ploidy.

We generated a series of isogenic haploid to tetraploid strains derived from the BY4748 strain background, all with the mating type MATa to eliminate the effects of the pheromone pathway (Extended Data 1A, Supplementary Table 1). Ploidy was confirmed by flow cytometry (Extended Data 1B). As expected, the volume of budding yeast cells increased linearly with ploidy (2,9), with median volumes of 48.0 fl for 1N, 82.9 fl for 2N, 146.6 fl for 3N and 181.7 fl for 4N (Extended Data 1C, D). Progression through the cell cycle was slightly reduced in tetraploids, as shown previously (7) (Extended Data 1E).

To assess the global proteome changes in strains of different ploidy, we used stable isotope labeling of amino acids in cell culture (SILAC) using an external spike-in standard consisting of a heavy lysine (Lys8)-labeled SILAC yeast standard, followed by liquid chromatography-tandem mass spectrometry (23). The heavy-labeled standard represented an equal protein mix from cells of different ploidy and was added to equal number of cells of different ploidy (Fig. 1A). The measurements provided quantitative information for 70% of all verified open reading frames in each strains with high correlation between independent experiments; this was also confirmed by principal component analysis (Extended Data 2A, B). 3109 protein groups quantified more than two times in three measurements were used to calculate the relative protein abundance in cells of each ploidy relative to the SILAC standard for normalization. This showed that the amount of proteins per cell increased with ploidy, but it did not scale linearly (Fig. 1B, 2N: 1.61, 3N: 2.31, 4N: 2.95). This “ploidy-specific protein scaling” (PSS) was validated by independent measurements of protein concentration and confirmed that the relative protein abundance per genome indeed decreases with increasing ploidy (Extended Data 2C, D).

**Figure 1.**
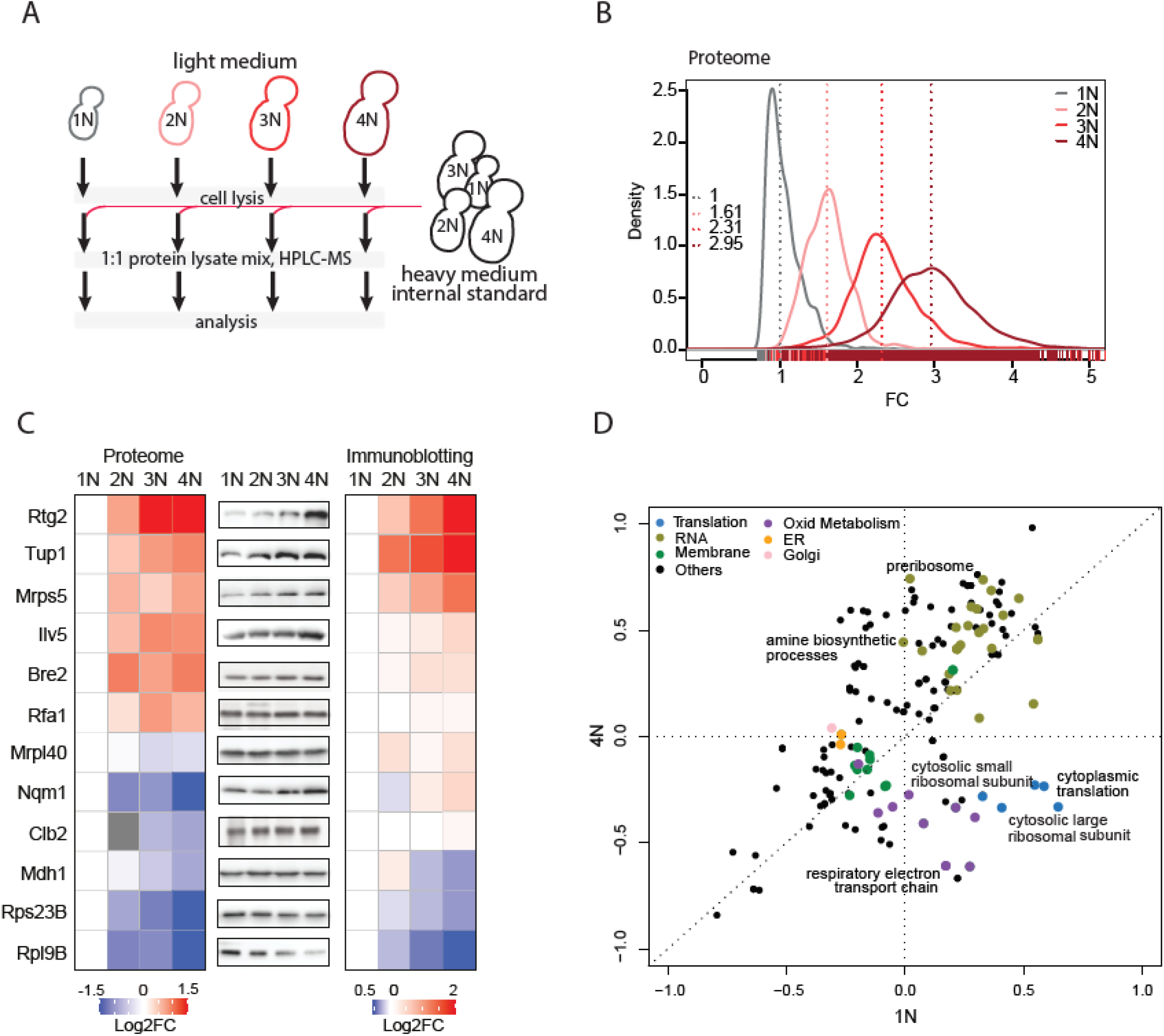
Proteome changes in response to increasing ploidy. **(A)** Schematic depiction of the strategy used for proteome analysis.**(B)** Proteome scaling with increasing ploidy. The values were normalized to the internal standard, with haploids set with a median at 1. 3109 were quantified in all ploidies. **(C)** Validation of the abundance changes of selected proteins. *Left*: relative proteome changes with ploidy normalized to 1N; *middle*: representative immunoblot of the selected candidates; *right*: Quantification of the protein abundance fold changes (FC) as determined by three biological replicates of immunoblotting, normalized to 1N. **(D)** Two-dimensional pathway enrichment analysis compares the proteome data of 1N and 4N cells. Each dot represents a pathway as determined by GO or KEGG databases. Only pathways significantly deregulated with respect to internal standard are depicted, Benjamini-Hochberg FDR <0.02. Related pathways are color-coded.

While most proteins followed the PSS trend, we found that several proteins were differentially regulated by ploidy (ploidy-dependent regulation, PDR). We calculated the log2 fold change relative to the SILAC standard and normalized the data by shifting the median of individual ploidies to 0 (Supplementary Data 1, the data can be visualized via a web-based application PloiDEx). This is not an artifact of SILAC or the normalization, as immunoblotting of 12 selected proteins matched well with the proteome results (Fig. 1C). To identify the differentially regulated proteins, we took two approaches: (1) we calculated 4N/1N ratio of protein abundance, and (2) classified proteins with a consistent abundance change across all ploidies (Extended Data 3A-D, for details see Methods). Among the proteins with ±2FC across ploidies, we found several upregulated cell wall integrity proteins (e.g. Cbk1, Prt1, Emw1, Fks1), whereas multiple mitochondrial (Isu2, Tma17, Tim13) and ribosomal (Rpl13A, Rpl31B) proteins were downregulated. Similar changes in protein abundance were observed also when yeast strains of a different genetic background were used, or when cultured in different media (Extended Data 3E-H). Moreover, PDR was not due to an increased volume of polyploid cells, as shown by analysis of two haploid mutants with altered cell size: cln3Δ that lacks a G1 cyclin, and a respiration deficient *rho^0^* mutant (Extended Data 4A, B). While the volume of these haploid mutants was comparable to that of wild type diploids and triploids (Extended Data 4B, C), the protein abundance of selected candidates did not correspond with the changes observed in polyploids (Extended Data 4D). Thus, ploidy changes induce differential regulation of several specific proteins.

Previous analyses of mRNA expression showed only marginal changes that could not explain the proteome changes described above (2,9,22). To confirm these findings in our strains, we used comparative differential transcription analysis (cDTA, (24)). mRNA was extracted from *S. cerevisiae* cells of different ploidy (1N, 2N, 3N, 4N) labeled with 4-thiouracil (4tU) and mixed it with mRNA of the distantly related haploid fission yeast *Schizosaccharomyces pombe* labeled with 4sU (4-thiouridin) as an internal standard (Extended Data 5A, (24)). This allowed us to obtain data on abundance changes for 5656 mRNAs. Comparison of mRNA abundance in cells of different ploidy showed similar ploidy-specific scaling as observed for protein abundance (Extended Data 5B). We then transformed the values to log2FC and normalized them by shifting the median of the distribution to 0. In agreement with previously published results (2,9,22), the expression of vast majority of mRNAs was not differentially expressed in polyploid cells (Data File 2, PloiDEx). Only 13 mRNAs change significantly with ploidy (±2FC), including factors involved in plasma membrane and cell wall synthesis (Extended Data 5C,D), in agreement with previous findings (9,22). Remarkably, qPCR analysis of selected factors whose protein levels were differentially regulated by ploidy confirmed that the changes were not present in the transcriptome (Extended Data 5E,F). Thus, the ploidy-dependent regulation of protein abundance occurs post-transcriptionally.

We next asked which pathways are differentially regulated in response to ploidy changes. Two-dimensional pathway enrichment comparison revealed striking ploidy-dependent down-regulation of pathways related to cytoplasmic ribosomes, translation, and mitochondrial respiration (Fig. 1D). Gene set enrichment analysis (GSEA) of the significantly deregulated proteins also confirmed that “ribosome biogenesis” and “cytoplasmic translation” were repressed with increasing ploidy, whereas vesicle trafficking and intracellular transport were activated (Extended Data 6A, B, Supplementary Data 3). Few pathways were differentially regulated at the mRNA level and there was no overlap with the proteome (Extended Data 6C, D), further confirming the low correlation between ploidy-dependent transcriptome and proteome changes. Proteome analysis showed that the respiratory electron transport chain was reduced. Consistently, the abundance of Rtg2, which plays a central role in retrograde signaling of mitochondrial dysfunction to the nucleus, was increased in cells of higher ploidy because of the stabilization of this protein (Fig. 1C, Extended Data 7A, B). Polyploid cells also proliferated poorly on media with non-fermentable carbon source or in the presence of the oxidant diamide (Extended Data 7C). This suggests ploidy-specific deregulation of mitochondrial function, in line with recent findings in pathogenic yeast *Candida albicans* (25).

Strikingly, “cytoplasmic translation” and “ribosome biogenesis” were strongly reduced with increasing ploidy (Fig. 1D, Extended Data 8A). We focused on this aspect, because it could explain the observed allometric PSS of protein content (Fig. 1B, Extended Data 2C, D). No mRNA changes could explain the altered abundance of ribosomal proteins, thus, we hypothesized that production of rRNA is reduced with increasing ploidy. Indeed, qRT-PCR revealed reduced abundance of 25S and 5.8 S rRNA in polyploid cells (Fig. 2A). We used pulse-labeling with puromycin, an amino acid analog that is incorporated into the nascent polypeptide chain and serves as a proxy for translational efficiency. This showed that the relative translation rate increased with ploidy when equal number of cells was loaded, but the increase was not linear, in agreement with proteome quantification; the puromycin incorporation appeared to be constant when equal amount of protein was loaded (Fig. 2B-D, Extended Data 8B).

**Figure 2.**
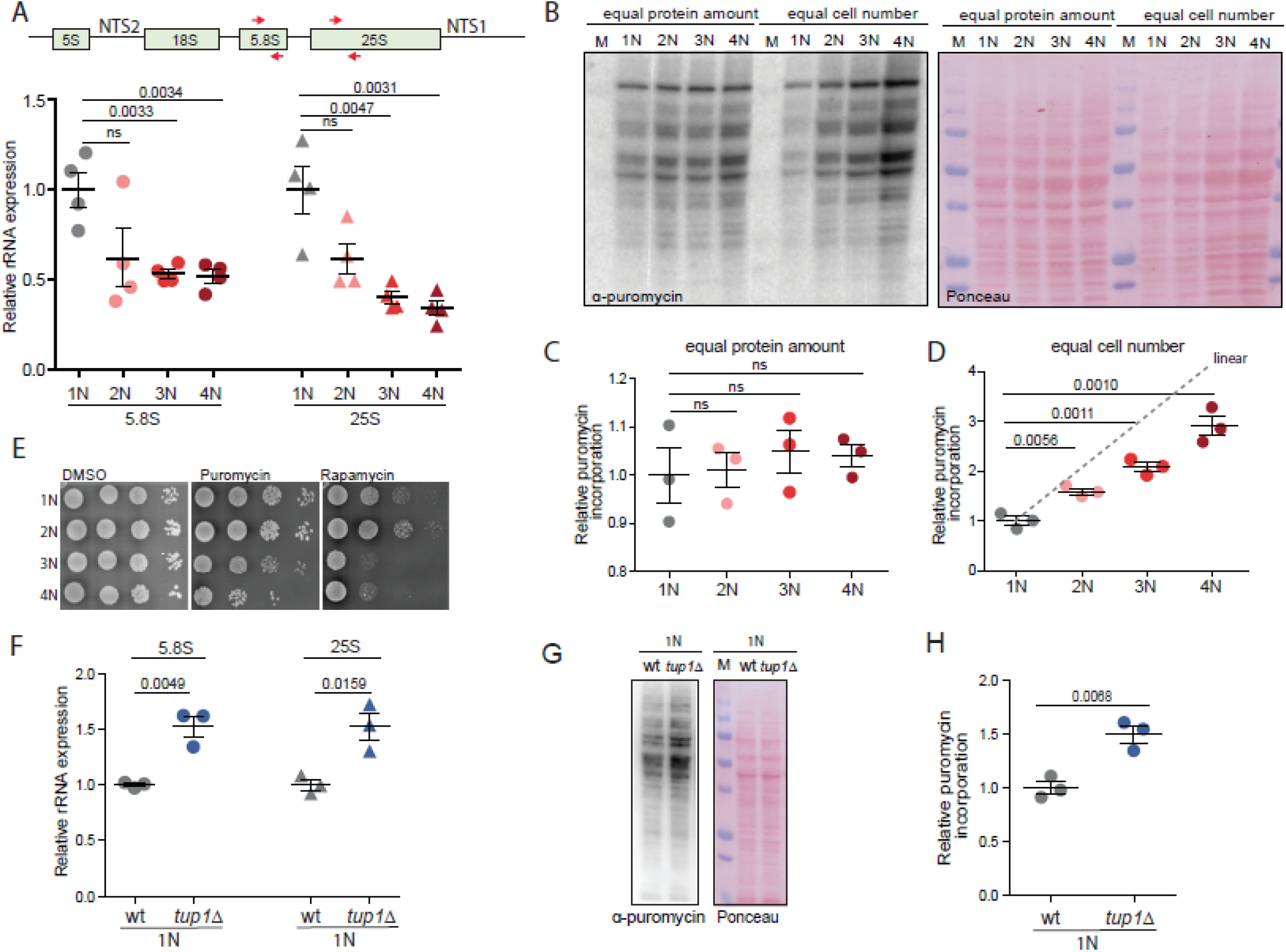
Translation and ribosome biogenesis are downregulated in cells with increased ploidy. **(A)** Quantification of rRNA abundance in cells of different ploidy. qRT-PCR was used in four independent experiments, the values were normalized to the expression of the housekeeping gene ACT1. Mean with SEM is shown. *Top*: schematics of the primers’ localization. **(B)** Puromycin incorporation in cells of different ploidy. Two types of loading were applied: *left*: equal amount of protein lysates were loaded, *right*: lysates from equal cell number was loaded. Ponceau staining was used as a loading control. **(C) (D)** Quantification of the relative puromycin incorporation calculated from three independent experiments; means with SEM are shown. **(E)** Sensitivity of cells of different ploidy to puromycin and rapamycin treatment. **(F)** Quantification of rRNA in haploid wt cells and in mutants lacking *TUP1*. The experiment was performed as in **(A)**. **(G)** Representative immunoblotting of puromycin incorporation in haploid wt and *tup1*Δ cells. **(H)** Quantification of puromycin incorporation from three independent biological replicates. Equal amount of protein lysate was loaded, Ponceau staining was used for the loading control. Means with SEM are shown.

The cells of higher ploidy were more sensitive to puromycin, an inhibitor of translation, as well as to rapamycin, an inhibitor of the master regulator of cellular metabolism mTOR, supporting the notion that the translational regulation is compromised via altered mTOR signaling (Fig. 2E).

What mechanism controls the reduced ribogenesis and translation in cells with higher ploidy? Because rRNA levels were reduced, we hypothesized that rDNA transcription decreases with increasing ploidy. We therefore examined all differentially regulated proteins for motives commonly found in repressors or for annotation indicating involvement in rRNA transcriptional repression. We identified Tup1, a protein whose human homolog Tle1 was previously shown to mediate rRNA repression (26). Accordingly, deletion of *TUP1* in 1N strains caused no changes in the abundance of several candidate mRNAs (Extended Data 8C), but lead to a significant increase in rRNA abundance and an increase in puromycin incorporation rates (Fig. 2F-H).

Ribosome biogenesis in eukaryotes is regulated via the mTOR pathway. Strikingly, treatment of haploid cells with rapamycin for 1 h increased Tup1 abundance to levels observed in tetraploids (Fig. 3A, B), clearly linking the regulation of Tup1 and rRNA gene expression to the mTOR pathway. In budding yeast, Tor1 phosphorylates the kinase Sch9, a yeast homolog of human P70-S6K, to promote ribosome biogenesis and protein synthesis. We asked whether Tor1-Sch9 regulates rDNA expression via Tup1. Haploid cells lacking Sch9 accumulated Tup1 independently of rapamycin treatment, suggesting that Tor1 negatively regulates Tup1 via Sch9 kinase (Fig. 3C). To test whether Tup1 is a downstream target of Sch9, we constructed a haploid strain with the analog-sensitive allele of Sch9 (Sch9-as, (27)). Inhibition with the ATP analog 1NM-PP1 increased the abundance of Tup1 and reduced its phosphorylation, as documented by altered migration in the Phos-TAG gel (Fig. 3D, Extended Data 8D). Finally, in haploid cells, loss of *TUP1* lead to constitutively high rRNA levels, while loss of *SCH9* reduced them (Fig. 3E). We asked whether the Tor1-Sch9 activity declines with increasing ploidy. Indeed, we observed decreased phosphorylation of Sch9 and Tup1, as judged by a reduced shift in migration on Phos-TAG gel, similar to mTOR inhibition with rapamycin (Fig. 3F, G, Extended Data 8E).Cyclohexamide shut off revealed that the stability of Tup1 increased in tetraploid cells, as well as in haploid cells lacking *SCH9* (Fig. 3H, I). We conclude that Tor1-Sch9 activity is reduced with increasing ploidy, which leads to accumulation of the negative regulator of rDNA expression Tup1. We propose that this novel signaling pathway is responsible for the ploidy-specific proteome scaling.

**Figure 3.**
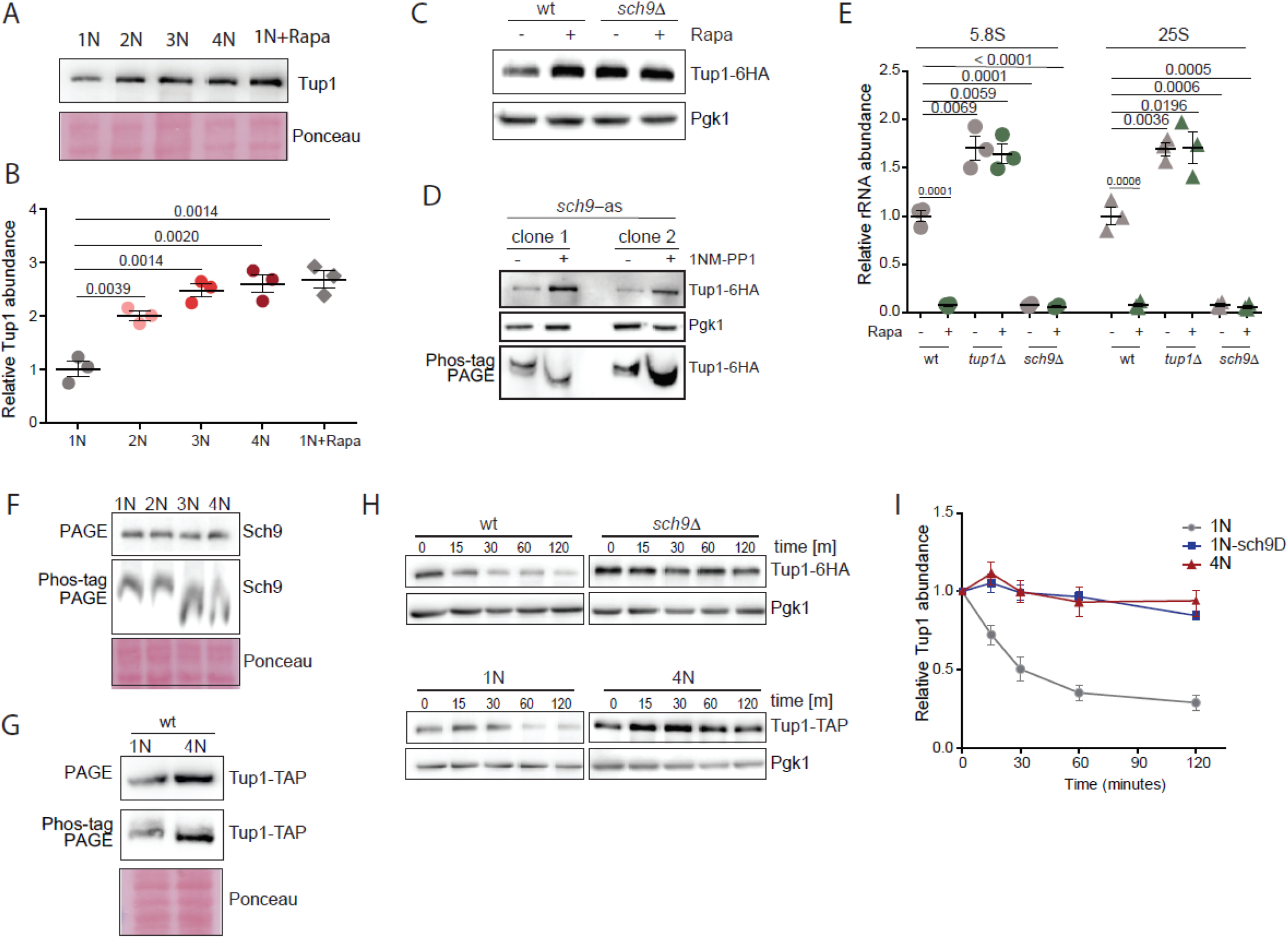
Regulation of rRNA expression in polyploid cells via Tor1-Sch9-Tup1 signaling pathway. **(A)** Relative abundance of Tup1 in cells of different ploidy compared to haploid cells treated with rapamycin. **(B)** Quantification of three independent experiments as in **(A)**. Mean with SEM is shown. **(C)** Representative immunoblot showing the abundance of Tup1 in wt and *sch9*Δ haploid cells treated with rapamycin. **(D)** Representative immunoblot showing the abundance of Tup1 in *sch9-as* haploid cells with and without the ATP analogue 1NM-PP1. Two different clones were tested. **(E)** qRT-PCR quantification of rRNA abundance in cells lacking Sch9 and Tup1, with and without rapamycin treatment (Rapa). Means and SEM of three independent experiments are shown. **(F)** Representative immunoblotting of Sch9 on PAGE and Phos-TAG gel in cells of different ploidy. **(G)** Representative immunoblotting of Tup1 on PAGE and Phos-TAG gel in cells of different ploidy. **(H)** Representative immunoblotting of Tup1 after cyclohexamide shut off. Different time points were collected and equal amount of lysate was loaded. **(I)** Quantification of three independent experiments from **(H)**. Means and SEM are shown.

We asked whether downregulation of translation and ribosome biogenesis occurs also in human tetraploid cells. To this end, we induced cytokinesis failure by treatment with the actin inhibitor dihydrocytochalasin D (DCD) in human diploid colorectal cancer cell line HCT116. This treatment induced formation of binucleated tetraploid cells (28). We also isolated a near-tetraploid cell line HPT2 (HCT116 Post Tetraploid) derived from a single HCT116 cell after WGD (Extended Data 9A)(17). Chromosome numbers and cell size increased with WGD as expected, as did the population heterogeneity (Fig. 4A, Extended Data 9B, C). While ribosomal protein expression was doubled in freshly formed binucleated tetraploids, four of six ribosomal proteins tested were reduced in HPT2 (Fig. 4B). Analysis of the previously obtained transcriptome of HPT2 confirmed that these changes were due to posttranscriptional regulation (17). Puromycin incorporation during translation was increased in binucleated tetraploids immediately after WGD, but was reduced in HPT2 below the diploid levels (Fig. 4C). This reduction was likely mediated via the mTORC1-S6K(Sch9)-Tle1(Tup1) pathway, as Tle1 abundance was elevated and mTORC1 activity, estimated by phosphorylation of S6K, was reduced in HPT2 but not in binucleated tetraploids (Fig. 4D-F, Extended Data 9D). Phosphorylation of the translation initiation factor 2, α subunit (eIF2 α) was not changed, suggesting that translational initiation was not affected by ploidy (Extended Data S9E). Finally, the expression of rRNA was reduced in the posttetraploid cells compared with the parental control (Fig. 4G). Thus, reduced mTORC1 activity and, in turn, reduced translation represents a conserved cellular response to increased ploidy and may be one of the major factors for the observed allometric scaling of protein levels with increased ploidy.

**Figure 4.**
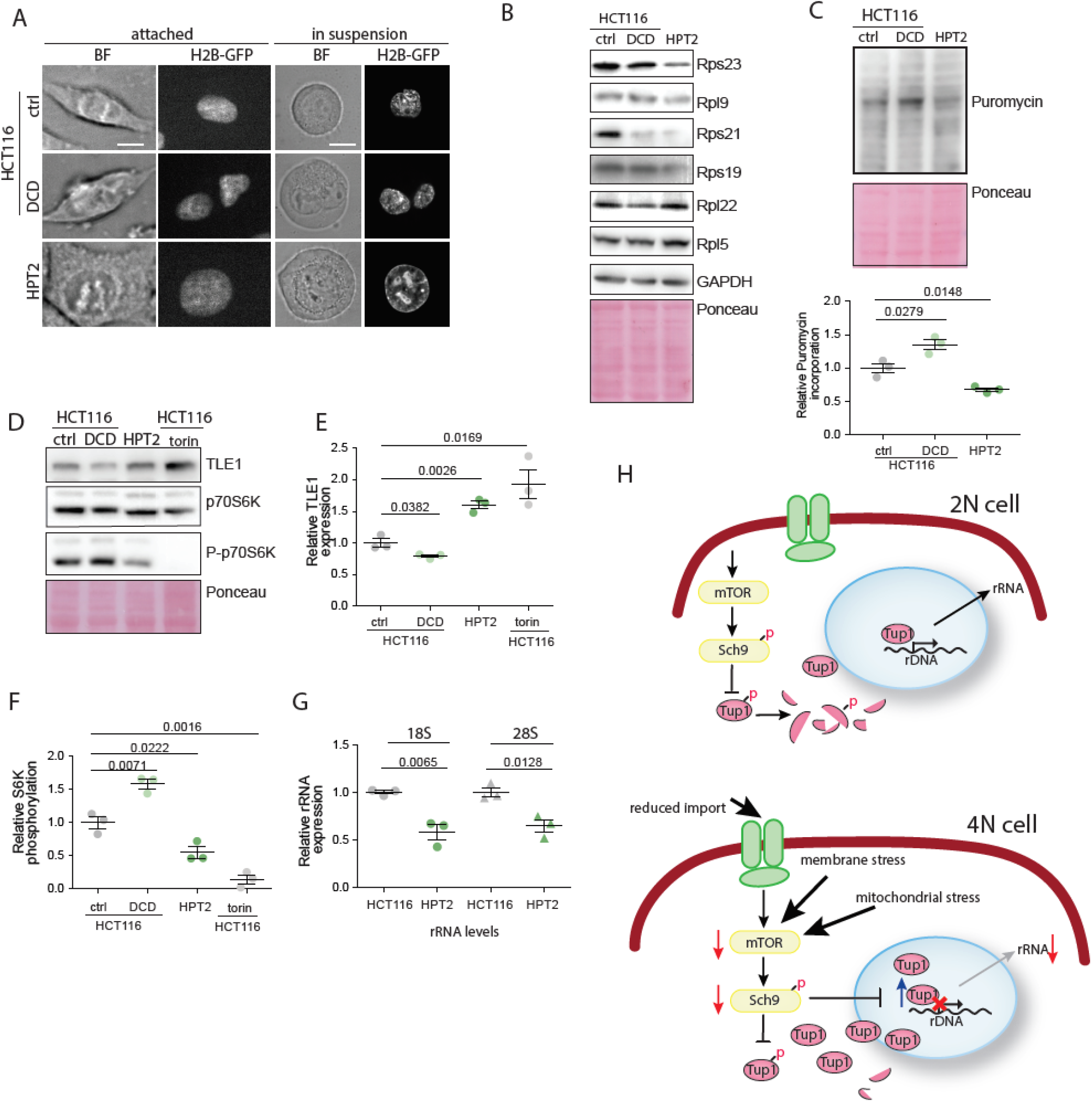
Downregulation of ribosomal proteins and translation in human near-tetraploid cells. **(A)** Cell size changes in response to altered ploidy. ctrl – untreated HCT116, DCD – HCT116 treated with dyhydrocytochalasin D to induce cytokinesis failure and subsequent tetraploidy; HPT2 – HCT116-derived near-tetraploid cell line. **(B)** Abundance of ribosomal proteins in response to altered ploidy. ctrl – untreated HCT116, DCD – HCT116 treated with dyhydrocytochalasin D to induce cytokinesis failure and subsequent tetraploidy; HPT2 – HCT116-derived near-tetraploid cell line. **(C)** Puromycin incorporation in diploid, newly made tetraploid HCT116 compared to near-tetraploid HPT2. Bottom: quantification of the relative puromycin incorporation. **(D)** Representative immunoblotting of Tle1 and S6K in human cells of different ploidy. **(E)** Quantification of Tle1, three independent experiments were analyzed. **(F)** Quantification of S6K from three independent experiments. **(G)** rRNA abundance in cells of different ploidy. Mean and SEM of three independent experiments is shown. **(H)** Schematic model of the translation regulation in polyploid cells.

This first proteome analysis in cells of different ploidy reveals striking ploidy-dependent proteome changes. First, we showed that protein abundance does not increase linearly with ploidy. Rather, it increases in accordance with allometric scaling theory where metabolic rate is predicted to scale with body size with an exponent close to ¾ (29,30), or, alternatively, with surface area with a scaling exponent of 2/3 (31). The observed PDS may have critical implication for cell physiology. Because the genome content and cellular volume increases linearly (2,9), allometric proteome scaling leads to cell dilution and a reduced ratio of DNA-binding proteins to DNA. Second, we show that many individual proteins are differentially regulated in response to ploidy. We call this phenomenon ploidy-dependent regulation – PDR. Our findings may help explain phenotypes of tetraploid cells that were difficult to reconcile with the minor changes observed in transcriptome and thus provide new hypotheses for mechanisms underlying ploidy-specific effects.

Strikingly, proteins related to ribosomes and translation were downregulated with increasing ploidy. The observed PDR of ribosome biosynthesis and translation also provided an explanation for the allometric proteome scaling, which we propose is regulated by the mTORC1-Sch9-Tup1 axes (Fig. 4H). Tup1 is a general transcription repressor that affects promoter accessibility (32). We found that Tup1 becomes stabilized by reduced Tor1-Sch9 activity in polyploids, resulting in reduced rRNA synthesis. The abundance of rRNA is limiting for ribosome biogenesis and thus the regulation of rRNA synthesis is central to overall ribosome synthesis. In agreement, the Tup1 homolog in humans, TLE1, previously shown to regulate rRNA expression (26), is also stabilized in tetraploid human cells.

What triggers reduced rDNA expression in polyploids? Ribosome biosynthesis is tightly regulated in response to environmental stress, and indeed stressors such as heat shock, osmotic shock, and nutrients deprivation reduce the expression of ribosomal proteins (33) and rRNAs (34). The reduced translation and rDNA expression could be a direct fitness cost of increased cell volume. The decreased surface-to-volume ratio may impair nutrient uptake, which in turn reduces mTOR activity and forces downregulation of protein synthesis.

Tetraploid yeast cells are sensitive to starvation (21) and often undergo ploidy reduction (3,8,19). Finally, stable tetraploid cells evolved for 1000 generations were smaller than the original cells, although they retained the tetraploid karyotype (35). These findings suggest that cell size might be a limiting factor causing strong selection to reduce ploidy or cell size, or both. Additionally, membrane stress due to abnormally large volume, and mitochondrial defect are known to alter mTOR activity and therefore may contribute to the observed PDR and PSS (36).

Another possibility is that downregulation of rDNA transcription could enhance adaptation to increased ploidy to prevent aberrant homologous recombination within the transcriptionally active rDNA repeats and R-loop formation. This hypothesis is alos supported by previous finding that the kinase Sch9 is required for the evolution of genomically stable yeast tetraploids (35). Genomic stability is impaired in cells with higher ploidy, and recombination rates increase significantly in budding yeast tetraploids (9). Because formation of R-loops threatens genome stability and increases recombination, frequent transcription of rDNA could be fatal to already compromised genomic stability in tetraploid cells.

Finally, an important observation is that PDR occurs largely on proteome level, via changes in protein stability. Proteome changes can occur rapidly and likely respond to more subtle environmental changes. Perhaps increased stress or prolonged *in vitro* evolution would be required to rewire the transcriptional patterns. Interestingly, many of the same genes and pathways identified here are altered at the transcriptional level after autopolyploidization in plants (37). Ribosomal and mitochondrial genes are also preferentially lost during evolution after whole genome duplication in Salmonidae (38). Together, these findings suggest conserved cellular changes triggered by increased ploidy that enable global metabolic adjustments to ensure survival of cells after whole genome doubling.

## Acknowledgments

We thank the members of Storchova lab, as well as Simen Rød Sandve and Johannes M. Herrmann for discussions and support. We thank Sara Bernhard for help with human tetraploid cells and Robbie Loewith for providing the Sch9-as coding plasmid. Funding: GY is a visiting scientist funded by Alexander von Humboldt Foundation (Georg Foster Stipend).

## Author Contributions

GY was involved in conceptualization, investigation, validation and funding acquisition, PM and MR were involved in investigation, formal analysis and data curation, AW contributed resources and investigation, DS and NK were involved in investigation, formal analysis and data curation, MM and PC were involved in conceptualization, supervision and funding acquisition, ZS was involved in conceptualization, writing, supervision and funding acquisition.

## Declaration of Interests

The authors declare no competing interests.

## Data and materials availability

The proteome and transcriptome data sets are available from public repositories, see details in Supplementary Materials.

## Methods

### Yeast media and culture

YP medium containing 1% yeast extract, 2% bacto-peptone, 2% dextrose, galactose or glycerol, respectively, were used. The synthetic drop-out (SC) medium consists of 5 g/L (NH4)2SO4, 2 g/L KH2PO4, 0.5 g/L MgSO4•7H2O, 0.1 g/L CaCl2•2H2O, 0.02 g/L FeSO4•7H2O, 0.01 g/L ZnSO4•7H2O, 0.005 g/L CuSO4•5H2O, 0.001 g/L MnCl2•4H2O, 1 g/L yeast extract, 10 g/L glucose, 0.5 mL/L 70% H2SO4200 μg/ml. G418 or 100 μg/ml ClonNat or 300 μg/ml Hygromycin were supplemented in YPD for drug resistance selection. For drop dilution, YPD pates were supplemented with 10 μg/ml Benomyl, 10 ng/ml Rapamycin, 500 ng/ml Tunicamycin 25 μM Puromycin, or 0.25 μM Diamide. For measurements of the protein half-life, the indicated strains were grown exponentially in YPD, then Cycloheximide (CHX) was added to a final concentration of 50 μg/mL to inhibit protein synthesis, the point of which was defined as time zero. Cell aliquots were collected at the indicated time points for protein extraction and western blotting. For experiments employing the sch9-as allele, 10 nM of 1NM-PP1 was used to inhibit the kinase activity of sch9-as.

### Yeast strain construction

To obtain yeast isogenic strain of different ploidies suitable for analysis, we used the strains of S288C genetic background BY4741 and BY4742. *LYS2* gene was deleted using xyz cassette to enable labeling with heavy lysine for SILAC experiments. BY4741 was transformed with a plasmid carrying the HO gene under the control of galactose-inducible promoter. Diploid strain was created by mating Mata and MATα haploid strains. Upon induction of HO expression, single diploid colonies that become mating-competent were selected and used for further mating to create triploid and tetraploid strain (Extended Data 1A). All strains were switch by HO expression to MATa mating type and the HO-expressing vector was removed by treatment with 5-FOA. Gene knockout strains were either retrieved from the available libraries deposit in public repository Euroscarf, or generated by homologous recombination using PCR products containing a drug cassette (kanMX6, Clonnat, hygMX) and 40 bp sequences flanking the target gene. Tagged-protein strains were either retrieved from the publicly available library (Euroscarf) or generated by integrating a cassette containing a protein tag and a drug resistance cassette at the C-terminus. PCR products were transformed into a haploid strain, or into diploid strains and the heterozygous diploids were sporulated and dissected to select for haploids with drug resistance. Subsequently, the procedure of polyploid construction described above was performed. For construction of strains with analog-sensitive sch9 allele, pRS414::sch9as (T492G) was used (a gift from Robbie Loewith, University of Geneva). The list of used strains is in Supplementary Table 1.

### Human cell line culture

HCT116 and the postetraploid derivative HPT2 were cultured in DMEM (Life Technologies) with 10% fetal bovine serum (Sigma-Aldrich) and 1%penicillin-streptomycin-glutamine (Life Technologies). Cells were incubated at 37°C, 5% CO2 and passaged twice a week using Trypsin-EDTA (0.25%) (Life Technologies). Cells were tested for mycoplasma contamination using the MycoAlert Mycoplasma Detection Kit (Lonza), according to the manufacturer’s instructions.

### Tetraploid formation in human cells

Cells were treated with 0.75 μM actin depolymerizing drug dihydrocytochalasin D (DCD, Sigma) for 18 hours. Subsequently, the drug was washed out 3x using prewarmed PBS. Cells were further cultured in drug-free medium for indicated time or immediately harvested for further experiments.

### Construction of post-tetraploids (WGD survivors)

HCT116 H2B-GFP were treated with 0.75 μM of the actomyosin inhibitor dihydrocytochalasin D (DCD, Sigma) for 18 hours. The cells were then washed, placed into a drug-free medium and subcloned by limiting dilution in 96-well plates (0.5 cell per well). After clone expansion, cells were harvested for flow cytometry to measure the DNA content. Subsequently, the genome was analyzed by SNP array analysis and multicolor fluorescent in situ hybridization to validate the tetraploidy. For further details see (17).

### RNA isolation

25 OD600 of yeast cells were collected from YPD exponential cultures, cell pellets were washed twice with ddH2O, then flash frozen in liquid nitrogen. 1 ml of trizol was added to frozen cell pellet and left on ice, cells then were resuspended and transferred to RNase-free screw cap Eppendorf containing 200 μl of acid washed nuclease free glass beads. Cells were disrupted by bead beating the tubes using the FastPrep-24™ 5G, 3 cycles of 20 secs at speed 6 m/s and 2 minutes in between each cycle. Add 200 μl chloroform, vortex 15 sec, then incubate at room temp for 5 min. Spin in Epp centrifuge, full speed, 5 min, 4°C. Recover supernatant to a fresh tube, and repeat trizol/ chloroform extraction with 1 ml trizol and 400 μl chloroform. Recover supernatant to a fresh tube and precipitate RNA by adding 0.5 ml of isopropanol and standing on ice for 15 minutes. Pellet by cold spin, wash twice with 1 ml 70% EtOH, Air dry for 10-20 min, then dissolve in an appropriate amount of RNase-free water depending on the pellet size. Measure the concentration and check the quality by Nano Drop, and agarose gel electrophoresis.

Total RNA was isolated from human cells (1 x 10^6^ cells) using the TRIZOL reagent according to the manufacturer’s instructions (ThermoFisher Scientific). RNA integrity for each sample was confirmed with by Nano Drop, and agarose gel electrophoresis.

### RT-qPCR

To assess the mRNA levels, total mRNA was isolated using a Qiagen mRNeasy mini kit according to manufacturer’s protocol. Next, reverse transcription using Anchored–oligo(dT) and Roche Transcriptor First Strand cDNA synthesis Kit (Cat no. 04 379 012 001) was performed to obtain cDNA. Quantitative PCR was performed using specific primers and SsoAdvanced Universal SYBR Green Supermix (Bio-Rad, USA). Melting curve analysis was performed to confirm the specificity of amplified product. Each sample was spiked with TATAA Universal RNA Spike II control (TATAA Biocenter AB, Sweden). mRNA expression of each sample was normalized to control housekeeping gene RPL30. The list of used primers is in Supplementary Table 3.

### Fluorescence microscopy

Exponentially growing yeast cells were imaged without fixation.

Human cells were seeded and treated when required in a glass-bottom 96-black well plate. The cells were then fixed using ice cold methanol, permeabilized with 3% Triton X 100 in PBS and blocked in blocking solution (5% Fetal Bovine Serum + 0.5% Triton X 100 + 1% Na3N in PBS). The DNA was stained DAPI or Vybrant DyeCycle™ Green for 1 hour in RT. Before imaging, cells were washed 4x with PBS. Imaging was carried out on a spinning disc system comprising of inverted Zeiss Observer.Z1 microscope, Plan Apochromat 63x magnification oil objective, 40x magnification air objective or 20x magnification air objective, epifluorescence X-Cite 120 Series lamp and lasers: 473, 561 and 660 nm (LaserStack, Intelligent Imaging Innovations, Inc., Göttingen, Germany), spinning disc head (Yokogawa, Hersching, Germany), CoolSNAP-HQ2 and CoolSNAP-EZ CCD cameras (Photometrics, Intelligent Imaging Innovations, Inc., Göttingen, Germany).

### Flow cytometry

For DNA content, 1 OD600 of yeast cells growing exponentially in YPD was collected and washed with ddH2O. Cells were fixed in 1 mL 70% (v/v) ethanol rotating at 4°C overnight. Fixed cells were then pelleted at 5000 rpm for 2 minutes and washed with ddH2O. Cells were subsequently resuspended in 500 μL FxCycle™ PI/RNase staining solution (Life Technologies, F10797), incubated at room temperature in the dark for 30 minutes and then stored at 4°C for 72 h or processed for next step. Samples were sonicated at 40% amplitude for 15 secs and run on an Attune™ Flow Cytometer. Data analyses was performed using the FlowJo™ software, version 10. Human cells were trypsinized and incubated in cold PBS supplemented with 5% fetal calf serum (Sigma-Aldrich; PBS-FACS). DNA was stained either by propidium iodide (PI); the cells were fixed in cold 70% ethanol added dropwise while vortexing, and incubated on ice for 30 minutes. Cells were centrifuged and pellets were washed twice with PBS-FACS. 50 μl RNase A solution (100 μg/ml in PBS) was added to the pellet, followed by staining with 400 μl PI solution (50 μg/ml in PBS) per million cells. Cells were incubated for 10’ at 25°C. For Hoechst staining, pellets were incubated in the dark with 10 mg/ml Hoechst 33358 for 15’ at 4°C. Data acquisition was performed using the ATTUNE NxT flow cytometer (ThermoFisher). Data analysis was performed using the FlowJo software. Gating strategy: An SSC-A/FSC-A gate was set in order to exclude cell debris, and an FSC-A/FSC-H gate was then set in order to exclude doublets.

### Cell size measurement

Cell volume of budding yeast was determined from microscopy bright field images of exponentially growing cultures using BudJ plugin according to (*Ferrezuelo et al., 2012*) To determine the cell volume of human cells, exponentially growing cells were harvested and washed with PBS twice. Forward scatter (FSC) was measured by ATTUNE NxT Flow Cytometer (ThermoFisher) and used as the indicator of cell size. Three biological replicated were examined and plotted.

### Cell growth for mass spectrometry

The cells were grown in SILAC synthetic drop out medium exponentially at a room temperature. The cell amount in each culture with different ploidy was counted using Burker chamber and equivalent amounts of cells of each ploidy were harvested and washed with PBS. To prepare the SuperSilac, the cells were grown in SILAC heavy medium, counted and mixed 1:1:1:1.

### Yeast sample preparation for proteomic analysis

Sample preparation was done as described as in (23). Briefly, cells were lysed in SDS lysis buffer (5% SDS, 100 mM dithiothreithol, 100 mM Tris pH 7.6), boiled for 5 min at 95°C and sonicated for 15 min (Bioruptor Sonicator, 20 kHz, 320 W, 60 s cycles). Insoluble remnants were removed by centrifugation at 16,000 g for 5 min and 140 μg clarified protein extract was transferred to a 30 kDa MW cut-off spin filter (Amicon Ultra 0.5mL Filter, Millipore). SDS was completely replaced by repeated washing with 8 M urea. Cysteines were then alkylated using excess amounts of iodoacetamide. Proteins were then proteolytically digested overnight using LysC endoprotease (1:50 w/w enzyme to protein). Peptides were eluted and desalted using C18 StageTips.

### Liquid chromatography coupled mass spectrometry

MS-based proteomic measurements were performed as in (23). Briefly, approximately 2 μg of desalted peptides were loaded and analyzed by linear 4h gradients. The LC system was equipped with an in-house made 50-cm, 75-μm inner diameter column slurry-packed into the tip with 1.9μm C18 beads (Dr. Maisch GmbH, Product Nr. r119.aq). Reverse phase chromatography was performed at 50°C with an EASY-nLC 1000 ultra-high-pressure system (Thermo Fisher Scientific) coupled to the Q Exactive mass spectrometer (Thermo Fisher Scientific) via a nano-electrospray source (Thermo Fisher Scientific). Peptides were separated by a linear gradient of buffer B up to 40% in 240 min for a 4-h gradient run with a flow rate of 250 nl/min. The Q Exactive was operated in the data-dependent mode with survey scans (MS resolution: 50,000 at m/z 400) followed by up to the top 10 MS2 method selecting 2 charges from the survey scan with an isolation window of 1.6 Th and fragmented by higher energy collisional dissociation with normalized collision energies of 25. Repeated sequencing was avoided using a dynamic exclusion list of the sequenced precursor masses for 40s.

### MS-Data Analysis

Raw files were analyzed by MaxQuant software version 1.6.3.3 (39) and searched against the *S. cerevisiae* Uniprot FASTA database (UniProt ID: UP000002311). Lysine-0 (light) and Lysine-8 (heavy) were used as SILAC labels. Cysteine carbamidomethylation was set as a fixed modification and N-terminal acetylation and methionine oxidation as variable modifications. LysC/P was set as protease and a maximum of two missed cleavages was accepted. False discovery rate (FDR) was set to 0.01 for peptides (minimum length of 7 amino acids) and proteins and was determined by searching against a generated reverse database. Peptide identification was performed with an allowed initial precursor mass deviation up to 7 ppm and an allowed fragment mass deviation of 20 ppm.

### Analysis of proteome data

Identified protein groups were filtered to remove contaminants, reverse hits and proteins identified by site only. SILAC light/heavy ratios were calculated and transformed to log2 scale. Next, Protein groups which were identified more than two times in at least one group of replicates were kept for further processing, resulting in a set of 3109 protein groups in total. To determine significance two-sample T-tests of 2N, 3N and 4N to 1N were performed (S0 = 0, permutation based Benjamini-Hochberg FDR threshold = 0.05). A “combined score” was calculated as the product of the q-values of all two sample tests. The median intensity of the replicates was calculated. The resulting data with statistics can be found in Supplementary Data 1.

Additional annotation (GOBP, GOCC) was added and 2D annotation enrichment analysis performed to identify significantly deregulated pathways (39). YeastEnrichR was used to perform gene set enrichment analysis of statistically significantly different values of all ploidies to haploid. The data is sorted based on the enrichR - combined score, which is a product of the p-value resulting from the Fisher exact test and the z-score of the deviation from the expected rank. Outliers were calculated twice based on the (SILAC L/H) ratio of ratios of each individual ploidy to 1N. First, the ratios of 4N to 1N were filtered for the outliers that show the overall strongest up or downregulation. Second, smoothed filtering was performed which additionally excluded values that show differences between consecutive ploidies with a FC >1 to remove values which spike between individual ploidies and keep only those that show a consistent trend across ploidies.

### PloiDEx - Supplementary plotting application

To allow users the plotting of our combined datasets PloiDEx (Ploidy-Dependent Expression), a suave.io based application has been written in the functional programming language F (4.5.2). The console application has been built in Visual Studio 2019 for the .NET Framework 4.6.1. It allows interactive plotting of both the proteome and transcriptome data, which have been normalized as described previously, in the user’s default browser using the graphing library plotly.js in the form of heatmaps and profile plots. For comparability the transcriptome has been matched to the proteome dataset. Filtering for outliers by a threshold and smoothing as described previously have been implemented. Additionally, the application allows the filtering of the dataset by GO cellular compartment or biological processes to plot the associated values from the database. For further information about the functionality of the application, used packages and libraries a Readme.pdf has been written, which together with the release and dependent packages is available at:

https://seafile.rlp.net/d/083c634e50d94e10834d/

### Dynamic transcriptome analysis (DTA)

We used the isogenic strains of different ploidies that were transformed with plasmid YEpEBI311 (2 μm, *LEU2*) carrying the human equilibrate nucleoside transporter hENT1. cDTA was performed as previously described (40). *S. cerevisiae* cells were grown in SD medium overnight, diluted to an OD_600_ of 0.1 the next day and grown up to a mid-log phase (OD_600_ of 0.8) and labeled with 4-thiouracil (4-tU, Sigma, 2M in DMSO) for 6 min at a final concentration of 5 mM. *Schizosaccharomyces pombe* cells were grown in YES medium (5 g/liter yeast extract; 30 g/liter glucose; supplements: 225 mg/liter adenine, histidine, leucine, uracil, and lysine hydrochloride) and labeled with 4sU (50 mM in ddH_2_O) for 6 min at a final concentration of 0.5 mM. A final concentration of 5 mM of 4tU was used. Cells were harvested via centrifugation and cell pellets re-suspended in RNAlater solution (Ambion/Applied Biosystems). The cell concentration was determined using a Cellometer N10 (Nexus) before flash freezing the cells in liquid nitrogen. Total RNA was extracted with the RiboPure-Yeast Kit (Ambion/Applied Biosystems), following the manufacturer’s protocol. Labeled RNA was chemically biotinylated and purified using strepatavidin-coated magnetic beads. Labeling of samples for array analysis was performed using the GeneChip 3IVT labeling assay (Affymetrix) with 100 ng input RNA. Samples were hybridized to GeneChip Yeast Genome 2.0 microarrays following the instructions from the supplier (Affymetrix).

### cDTA data analysis

Analysis of cDTA data was carried out as described (40) using R/Bioconductor and the DTA package. Briefly, array probes that cross-hybridized between *S. pombe* and *S. cerevisiae* were removed from the analysis. The labeling bias was performed as described (24) and proportional rescaling of the *S. cerevisiae* microarray intensities to the internal *S. pombe* standard was performed to obtain global expression fold changes (40). Differential gene expression analysis was performed using the R Bioconductor package “Limma” and multiple testing correction was done using false discovery rates.

### Protein isolation from budding yeast and human cells

Exponentially growing yeast cells were harvested, resuspended in 100 μl lysis buffer and incubated 10 min on ice. 40 μl 100% TCA was added and incubated for 10 min on ice. Precipitated proteins were spun down for 10 min at 4°C, 13 000. The pellet was washed with 1 ml ice-cold acetone, dried at 50°C for 5-20 min and resuspended in 50 μl 2x Laemmli buffer. Protein lysates were boiled for 5 min at 96°C. Pelleted human cells were lysed in RIPA buffer with protease inhibitor cocktail (Pefabloc SC, Roth, Karlsruhe, Germany), then sonicated by ultrasound in a water bath for 15 min. Cell lysate was spun down at 13600 rpm for 10 minutes at 4°C on a table-top microcentrifuge (Eppendorf, Hamburg, Germany). 1 μl was used to determine protein concentration using Bradford dye at 595 nm wavelength. Subsequently, the lysates were mixed with 4x Lämmli buffer with 2.5%ß-mercaptoethanol and boiled at 95°C for 5 min.

### Immunoblotting

Prepped cell lysates were separated by SDS-PAGE using 10% or 12.5% gels. Protein size was estimated using the PrecisionPlus All Blue protein marker (Bio-Rad, USA). Gels were incubated in Bjerrum Schafer-Nielsen transfer buffer and proteins were transferred to a water-activated nitrocellulose membranes (Amersham Protran Premium 0.45 NC, GE Healthcare Life Sciences, Sunnyvale, USA) using semi-dry transfer (Trans-Blot® Turbo™, Bio-Rad, USA). Membranes were stained in Ponceau solution for 5 min and scanned to be used as a loading control. Next, membranes were blocked in 5% - 10% skim milk in TBS-T (Fluka, Taufkirchen, Germany) for 1 hour in RT. After blocking, membranes were incubated in respective primary antibodies diluted in 1% Bovine Serum Albumin (BSA) or 5% skim milk overnight at 4°C with gentle agitation. Further, the membranes were rinsed 3 x 5 minutes with TBS-T, incubated 1 hour in RT with HRP-conjugated secondary antibodies (RD Systems), and followed by rinsing 3x 5 minutes with TBS-T. Chemiluminescence was detected using ECLplus kit (GE Healthcare, Amersham™) and Azure c500 system (Azure Biosystems, Dublin, USA). Protein band quantification was carried out using ImageJ (National Institutes of Health, http://rsb.info.nih.gov/ij/). List of used antibodies is in Supplementary Table 2.

### Phos-tagTM SDS-PAGE

Phospho-affinity gel electrophoresis for mobility shift detection of phosphorylated proteins was performed using Phos-tag acrylamide 4.5% (w/v) running gels polymerized with 25 μM Phos-tag acrylamide (FUJIFILM Wako Pure Chemical Corporation) and 50 μM MnCl2. Gel running and transfer conditions were optimized according to the manufacturer’s protocol.

### Puromycin incorporation

Yeast cells were grown exponentially in YPD media then puromycin was added to the culture at a final concentration of 10 μM for 15 minutes shaking at 30°C, then 5 OD_600_ of cells were collected, washed twice with ddH_2_O and prepared for western blot.

### Data availability and accession

Transcriptome data have been deposited in the Gene Expression Omnibus database under accession code GSE162513.

The mass spectrometry proteomics data have been deposited to the ProteomeXchange Consortium via the PRIDE partner repository with the dataset identifier PXD022605.

## Supplementary Information is available for this paper

Supplementary Figures 1-9

Supplementary Tables 1-4

Supplementary Data Files 1-3

## Notes

### Competing Interest Statement

The authors have declared no competing interest.

### Summary of Updates

Spelling error in authors' list was corrected.

