## Supplemental Information for "Scaling of cellular proteome with ploidy" (from another version of this preprint: version 1, 2021-05-07)

**Extended Data**

**This PDF file includes:**

Extended Data 1 to 9
Supplementary Tables 1 to 4
Captions for Supplementary Data Files 1, 2 and 3

**Other Supplementary Materials for this manuscript include**

Supplementary Data Files 1, 2 and 3 (excel tables)

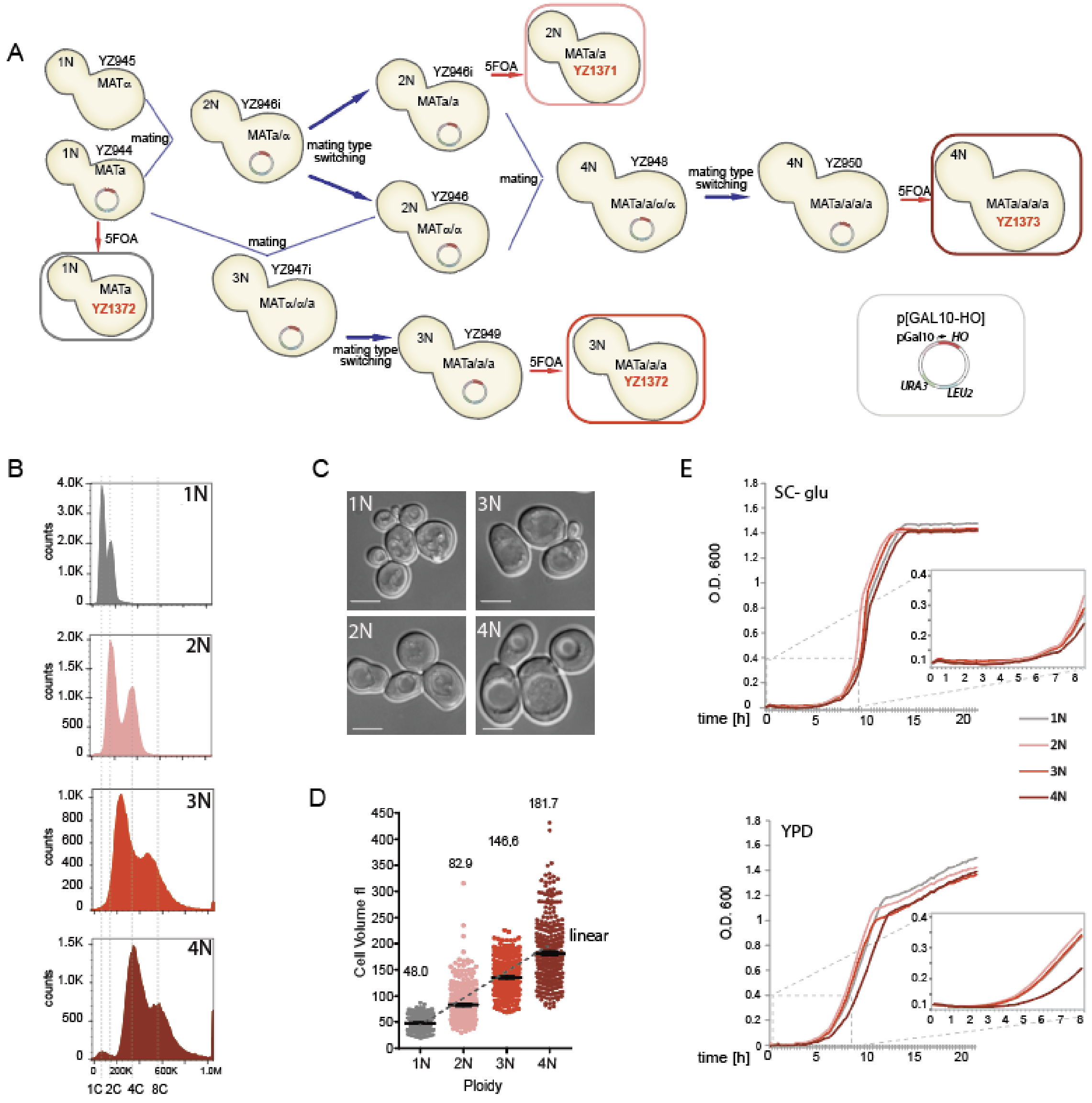

**Extended Data 1. Cell volume and proliferation changes in cells of different ploidy**

(A) Schematic depiction of the construction of isogenic yeast strains that differ only in their ploidy. The plasmid p[Gal10-HO] was used for the inducible mating type switching. The marked strains were used for the analysis. Further details can be found in Material and Methods. Bottom right: Schematics of the plasmid carrying the inducible HO endonuclease. (B) Flow cytometry analysis to determine the DNA content of the used strains. (C) Representative images of yeast cells of different ploidy. Scale bar 5mm. (D) Quantification of the cell volume of yeast cells of different ploidy. 400 asynchronous cells were analyzed in four independent experiments, only small budded cells were included in the analysis and the volume of the mother cell was measured. Dashed line marks linear scaling. Top values: Mean volume. (E) Proliferation of yeast cells of different ploidy in fully supplemented synthetic medium with dextrose as the carbon source (SC-glu) and in a complete medium with dextrose (YPD) at 30° C.

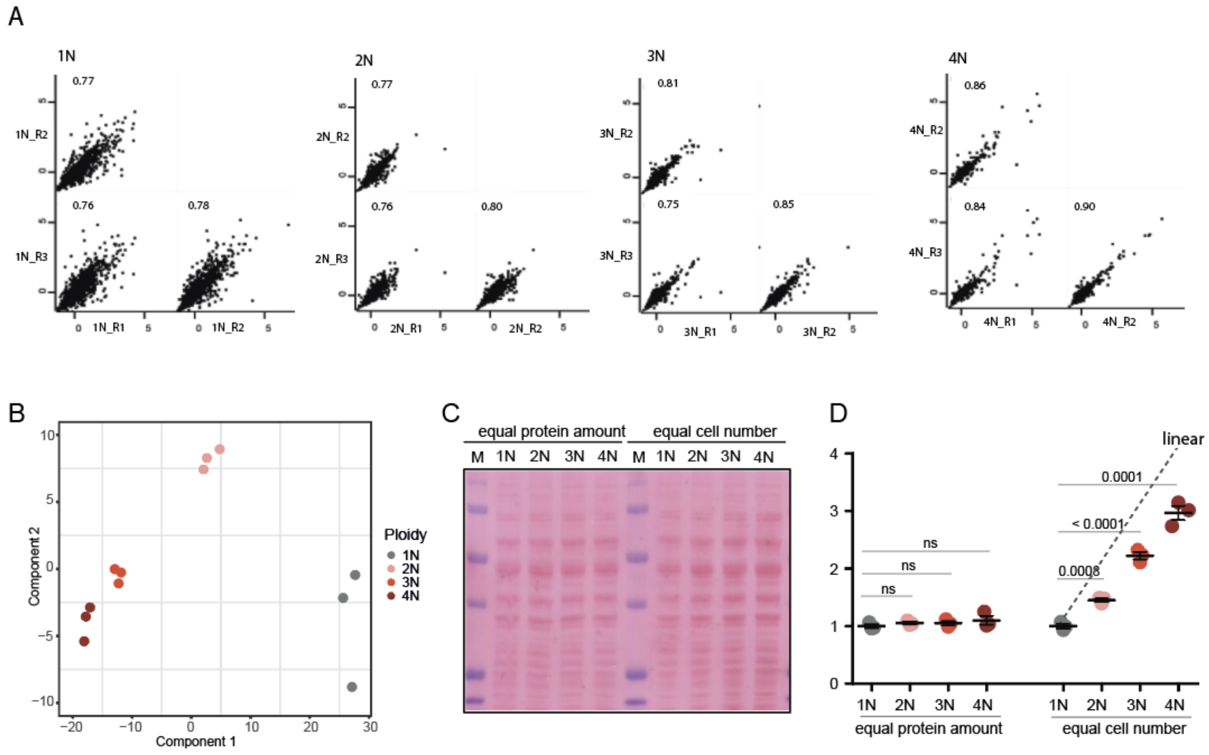


**Extended Data 2. Proteome analysis in cells of different ploidy**

(A) Correlation between runs of the MS/MS analysis. (B) Principal component analysis of the individual MS/MS measurements. (C) Representative Ponceau staining of the total protein amount in cell of different ploidy when equal protein amount is loaded (left) and when proteins isolated from equal cell numbers are loaded (right). (D) Quantification of the protein levels in (C). Three independent experiments were performed. Dashed line marks linear scaling.

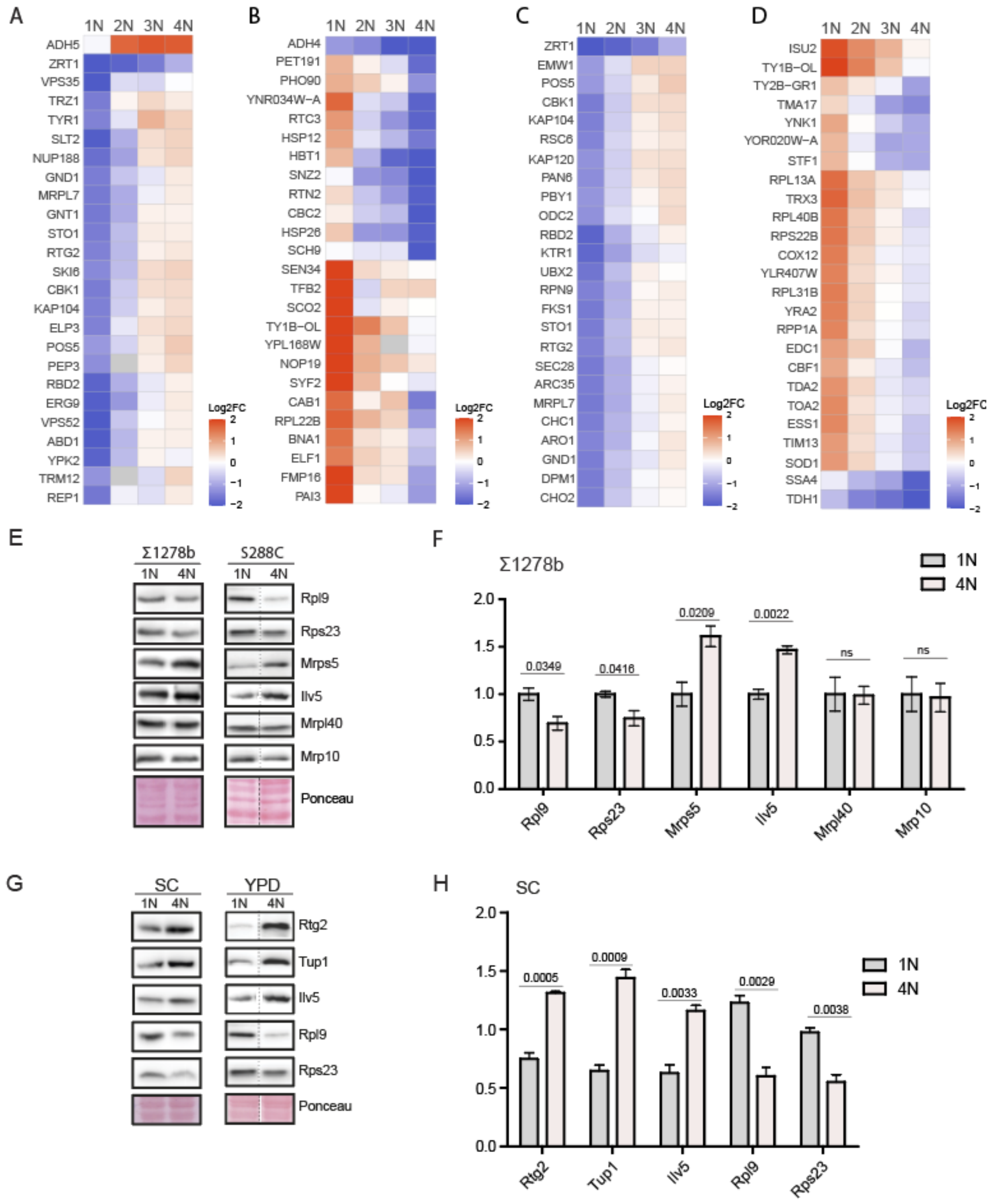


**Extended Data 3. Proteins differentially regulated with ploidy**

(A) Top 25 proteins upregulated in tetraploids (4N) compared to haploids (1N). (B) Top 25 proteins downregulated in 4N compared to 1N. (C) Top 25 proteins upregulated across ploidies. (D) Top 25 proteins downregulated across ploidies. (E) Representative immunoblot of selected proteins in 1N and 4N cells derived from Σ1278b background (left) and from S288C background. (F) Quantification of the immunoblotting in Σ1278b. Means with SEM of three independent experiments are shown. (G) Representative immunoblot of selected proteins in 1N and 4 N cells grown in synthetic (left) and rich media. (H) Quantification of the immunoblotting in SC. Means with SEM of three independent experiments are shown.

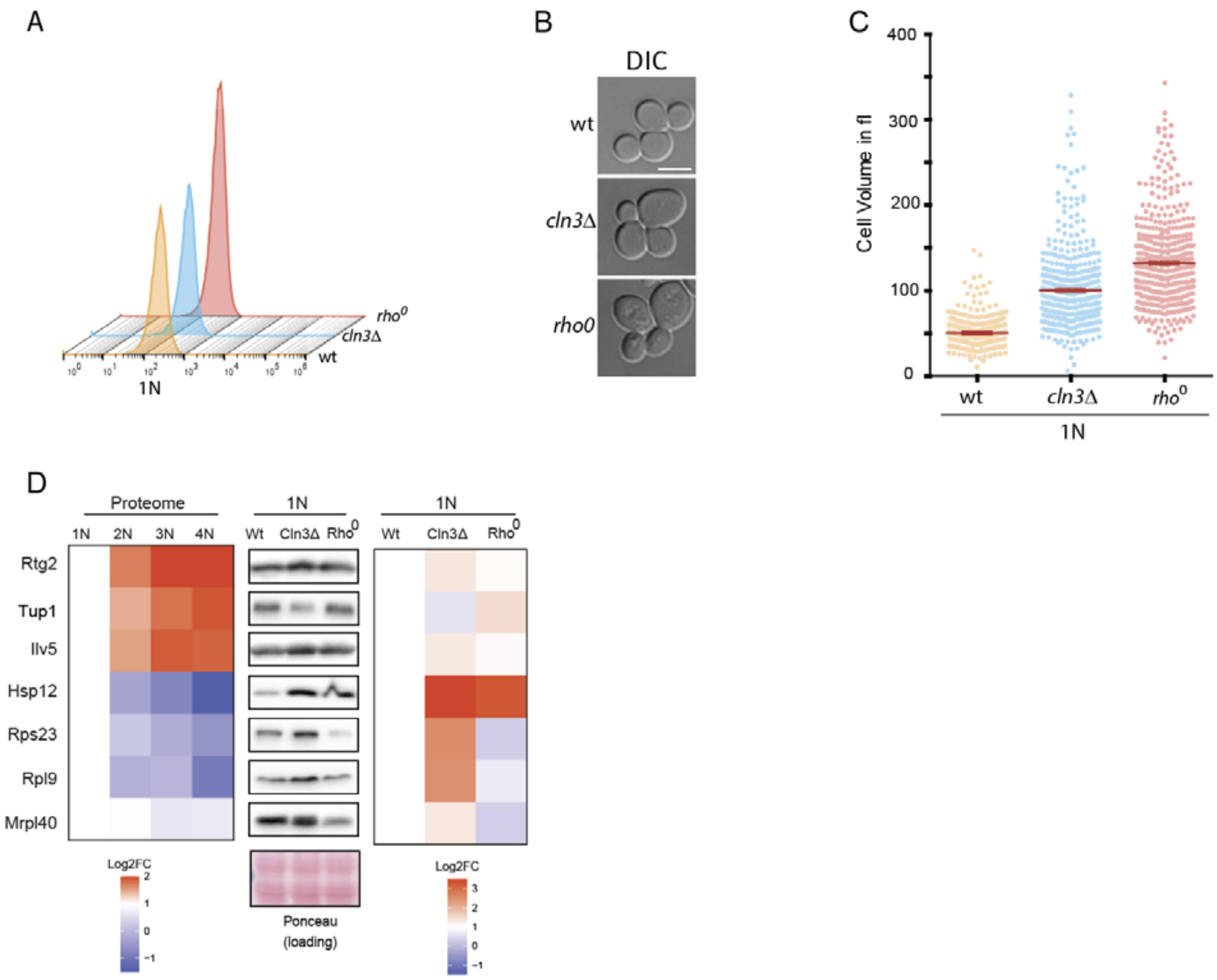


**Extended Data 4 Cell volume-dependent gene expression changes differ from changes due to increased ploidy**

(A) Histogram of the DNA content in the used mutant strains as measured by flow cytometry. (B) Representative images of the analyzed yeast mutants. Scale bar 5 mm. (C) Quantification of the cell volume of the yeast mutants. Cell volume of >400 cells was calculated as indicated in the Materials and Methods. The average cell volumes of the indicated strains were 50.43, 100.3, and 132.4 fl, respectively. (D) Representative immunoblot and quantification of protein levels based on immunoblotting of whole cell lysates of the respective haploid mutant strains and their comparison to yeast strains of different ploidy (normalized to 1N).

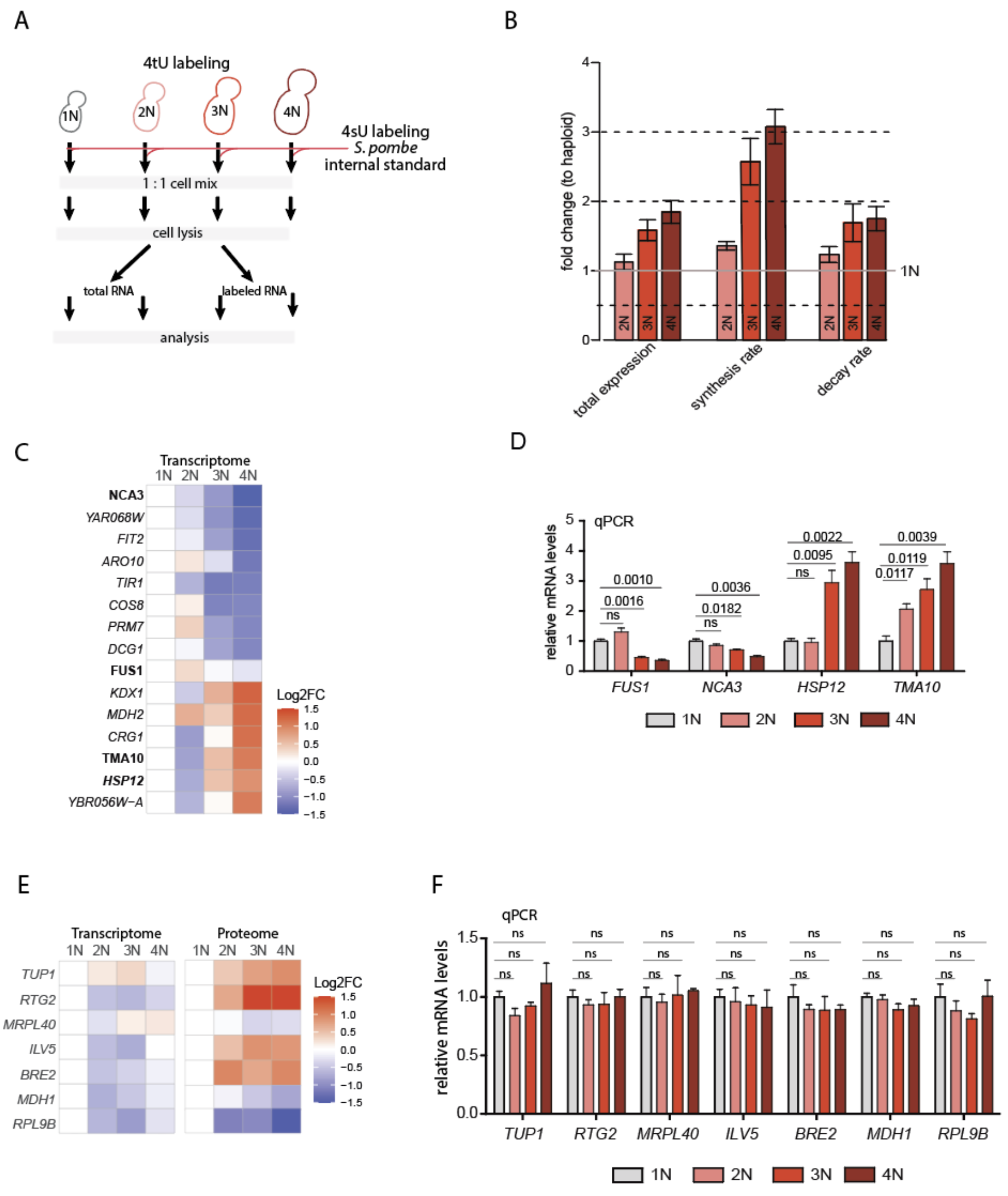


**Extended Data 5 Transcriptome changes in cell with altered ploidy**

(A) Schematic depiction of the strategy used for transcriptome analysis. (B) Dynamic mRNA abundance changes in strains of different ploidy compared to 1N strain. (C) Heat map of the most differentially regulated transcripts normalized to 1N. Transcripts marked in bold were validated by qRT-PCR. (D). Quantification of mRNA levels of selected differentially regulated candidates in cells of different ploidy from (B), normalized 1N. Mean values and SD of three independent experiments are shown. (E) Heat map of selected candidates' mRNAs that were not differentially regulated by ploidy at a transcriptome level, but changed on proteome level. (F) Quantification of mRNA abundance by qRT-PCR of the candidates from (D), normalized to 1N. Means and SD of three independent experiments are shown.

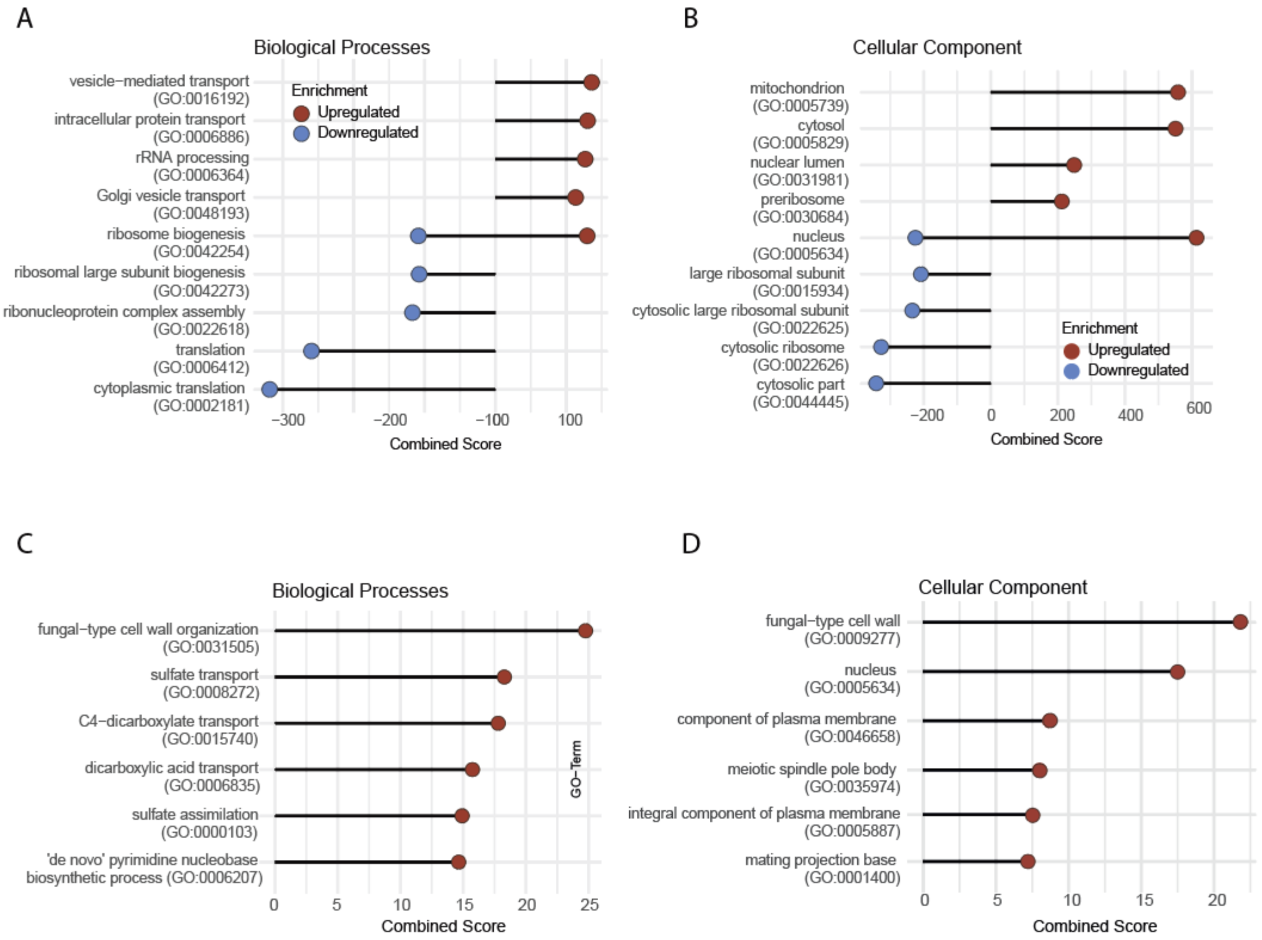


**Extended Data 6 Differentially regulated pathways**
(A) Differentially regulated Biological Process and (B) Cellular Components identified by Gene Set Enrichment Analysis of proteome. (C) Differentially regulated Biological Processes and (D) Cellular Components identified by Gene Set Enrichment Analysis of mRNAs. No downregulated pathways were identified by transcriptome.

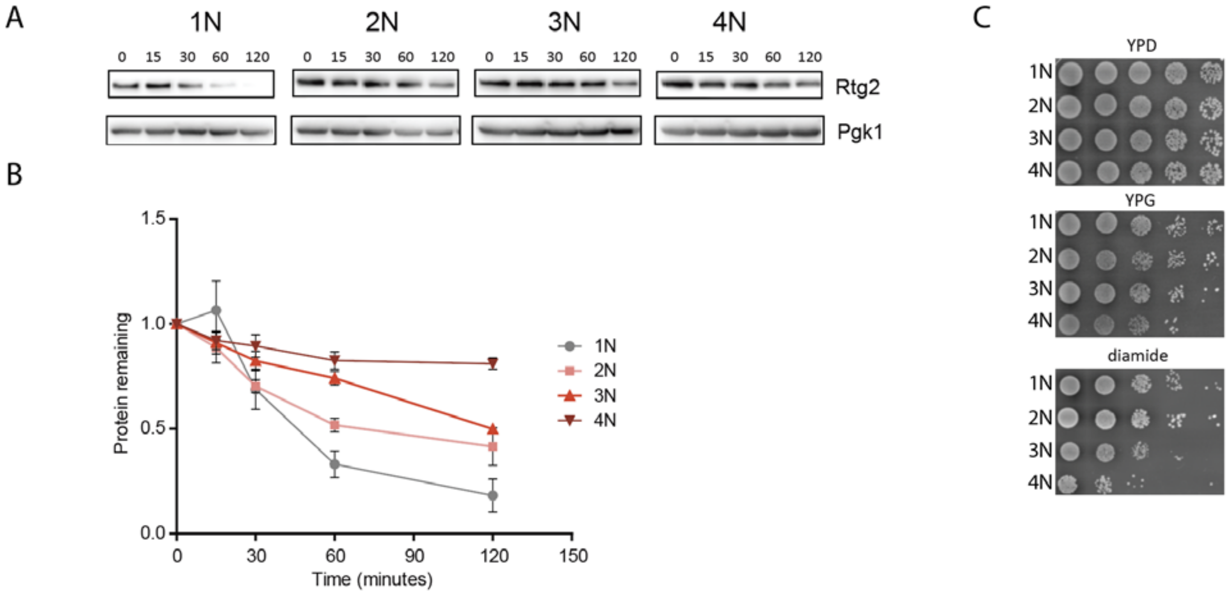


**Extended Data 7 Deregulation of mitochondrial metabolism in cells with higher ploidy**
(A) Representative immunoblot of the protein abundace of Rtg2 after a cyclohexamide shut off at respective time points in cells of different ploidy. (B) Quantification of the Rtg2 abundance after cyclohexamide shut off. Means and SD of three independent experiments are shown. (C) Reduced proliferation of cells with different ploidy on non-fermentable medium and in presence of oxidazing agent diamide.

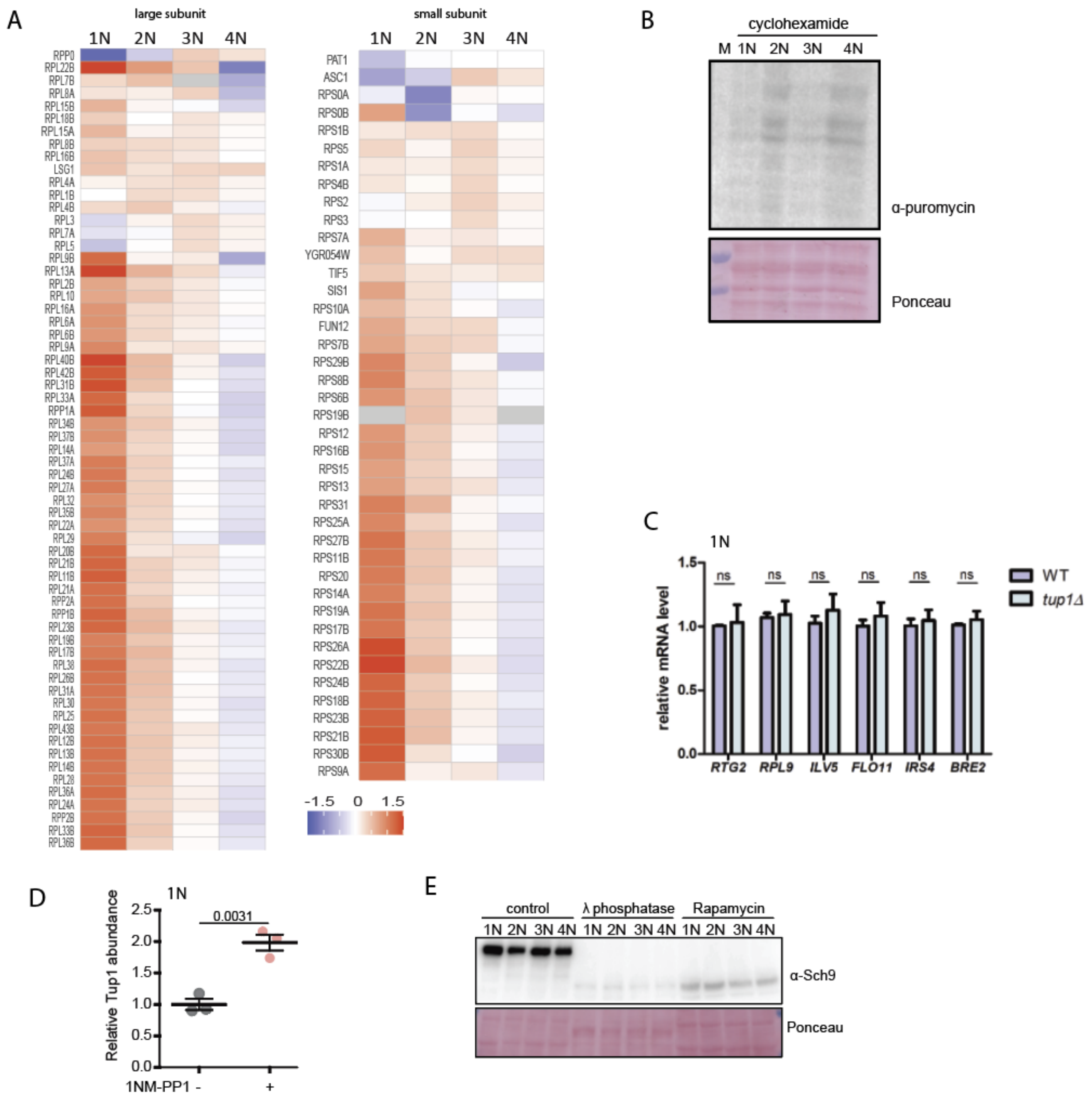


**Extended Data 8 Translation downregulation in cells with increased ploidy**
(A) Abundance of ribosomal proteins in cells of different ploidy. (B) Treatment with cyclohexamide abolishes puromycin incorporation. Control to Fig. 2B. (C) Abundance of mRNAs in haploid wild type (WT) and *tup1Δ* strain quantified by RT PCR.(D) Relative abundance of Tup1 in haploid cells carrying the analog-sensitive *SCH9* allele with and without 1NM-PP1 treatment. (W) Abundance of Sch9 and its phosphorylation in cells of different ploidy and upon rapamycin and phosphatase treatments.

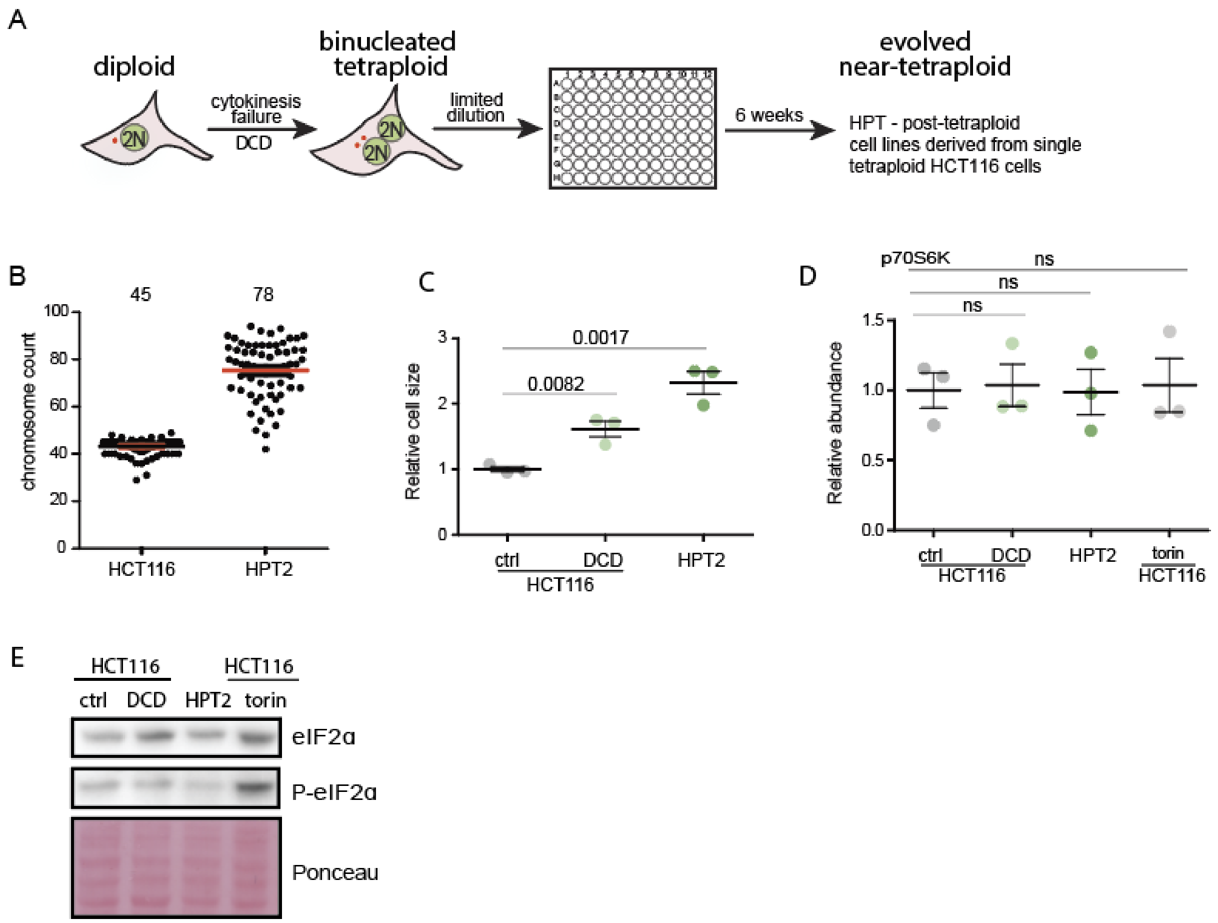


**Extended Data 9 Human tetraploid and near-tetraploid cell lines**
(A) Schematic depiction of the construction of post-tetraploid cell line HPT2. (B) Chromosome count in parental HPT116 and in posttetraploid HPT2 clone. (C) Relative cell volume evaluated by flow cytometry in diploid HCT116 cells (ctrl), in binucleated tetraploids shortly after DCD tratment (DCD) and in the posttetraploid cells (HPT2). (D) Relative abundance of p70-S6K. Mean and SEM are shown in the plots C, D; three biologically independent experiments were performed. (E) Representative immunoblot of eIF2α and P-eIF2α in diploid and tetraploid cells.

### Supplementary Table 1. A list of used yeast strains

All laboratory strains were constructed in the laboratory for this work, unless otherwise stated.

| Name | Genotype | Plasmid | Remarks |
|---|---|---|---|
| BY4741 | *MATa his3Δ1 leu2Δ0 met15Δ0 ura3Δ0* | | S288C Euroscarf |
| BY4742 (YZ945) | *MATα his3Δ1 leu2Δ0 lys2Δ0 ura3Δ0* | | S288C Euroscarf |
| YZ944 | *MATa his3Δ1 leu2Δ0 met15Δ0 ura3Δ0 lys2Δ* | PB1650 [*LEU2, URA3, pGal-HO*] | |
| YZ946i | *MATa/α his3Δ1/his3Δ1 leu2Δ0/leu2Δ0 met15Δ0/MET15 ura3Δ0/ura3Δ0 lys2Δ/lys2Δ* | PB1650 [*LEU2, URA3, pGal-HO*] | |
| YZ946 | *MATα/α his3Δ1/his3Δ1 leu2Δ0/leu2Δ0 met15Δ0/MET15 ura3Δ0/ura3Δ0 lys2Δ/lys2Δ* | PB1650 [*LEU2, URA3, pGal-HO*] | |
| YZ947 | *MATa/a his3Δ1/his3Δ1 leu2Δ0/leu2Δ0 met15Δ0/MET15 ura3Δ0/ura3Δ0 lys2Δ/lys2Δ* | PB1650 [*LEU2, URA3, pGal-HO*] | |
| YZ947i | *MATa/a/α his3Δ1/his3Δ1/his3Δ1 leu2Δ0/leu2Δ0/leu2Δ0 met15Δ0/MET15/MET15 ura3Δ0/ura3Δ0/ura3Δ0 lys2Δ/lys2Δ/lys2Δ* | PB1650 [*LEU2, URA3, pGal-HO*] | |
| YZ948 | *MATa/a/α/α his3Δ1/his3Δ1/his3Δ1/his3Δ1 leu2Δ0/leu2Δ0/leu2Δ0 /leu2Δ0 met15Δ0/MET15/MET15/met15Δ0 ura3Δ0/ura3Δ0/ura3Δ0/ura3Δ0 lys2Δ/lys2Δ/lys2Δ/lys2Δ* | PB1650 [*LEU2, URA3, pGal-HO*] | |
| YZ949 | *MATa/a/a his3Δ1/his3Δ1/his3Δ1 leu2Δ0/leu2Δ0/leu2Δ0 met15Δ0/MET15/MET15 ura3Δ0/ura3Δ0/ura3Δ0 lys2Δ/lys2Δ/lys2Δ* | PB1650 [*LEU2, URA3, pGal* | |
| YZ950 | *MATa/a/a/a his3Δ1/his3Δ1/his3Δ1/his3Δ1 leu2Δ0/leu2Δ0/leu2Δ0 /leu2Δ0 met15Δ0/MET15/MET15/met15Δ0 ura3Δ0/ura3Δ0/ura3Δ0/ura3Δ0 lys2Δ/lys2Δ/lys2Δ/lys2Δ* | PB1650 [*LEU2, URA3, pGal-HO*] | |
| YZ1375 | As YZ944 | | Used for SILAC |
| YZ1371 | As YZ947 | | Used for SILAC |
| YZ1372 | As YZ949 | | Used for SILAC |
| YZ1374 | As YZ950 | | Used for SILAC |
| YZ1535 | *MATa/leu2Δ/ura3Δ/his3Δ/met15Δ/NQM1::TAP* | | |
| YZ1547 | *MATa/leu2Δ/ura3Δ/his3Δ/met15Δ/BRE2::TAP* | | |
| YZ1554 | *MATa/leu2Δ/ura3Δ/his3Δ/met15Δ/TUP1::TAP* | | |
| YZ1556 | *MATa/leu2Δ/ura3Δ/his3Δ/met15Δ/RTG2::TAP* | | |
| YZ1562 | *MATa/a/leu2Δ/ leu2Δ/ura3Δ/ ura3Δ/his3Δ/ his3Δ/met15Δ/met15Δ/NQM1::TAP/NQM1::TAP* | | |
| YZ1566 | *MATa/a/leu2Δ/ leu2Δ/ura3Δ/ ura3Δ/his3Δ/ his3Δ/met15Δ/met15Δ/BRE2::TAP/BRE2::TAP* | | |

| | | | |
|---|---|---|---|
| YZ1602 | *MATa/a/leu2Δ/ leu2Δ/ura3Δ/ ura3Δ/his3Δ/ his3Δ/met15Δ/met15Δ/RTG2::TAP/RTG2::TAP* | | |
| YZ1603 | *MATa/a/leu2Δ/ leu2Δ/ura3Δ/ ura3Δ/his3Δ/ his3Δ/met15Δ/met15Δ/TUP1::TAP/TUP1::TAP* | | |
| YZ1605 | *MATa/a/a/leu2Δ/ leu2Δ/leu2Δ/ura3Δ/ura3Δ/ ura3Δ/his3Δ/ his3Δ/ his3Δ/met15Δ/met15Δ/met15Δ/NQM1::TAP/NQM1::TAP/NQM1::TA P* | | |
| YZ1607 | *MATa/a/a/leu2Δ/ leu2Δ/leu2Δ/ura3Δ/ura3Δ/ ura3Δ/his3Δ/ his3Δ/ his3Δ/met15Δ/met15Δ/ met15Δ/BRE2::TAP/BRE2::TAP/BRE2::TAP* | | |
| YZ1617 | *MATa/a/a/leu2Δ/ leu2Δ/leu2Δ/ura3Δ/ura3Δ/ ura3Δ/his3Δ/ his3Δ/ his3Δ/met15Δ/met15Δ/ met15Δ/RTG2::TAP/RTG2::TAP/RTG2::TAP* | | |
| YZ1614 | *MATa/a/a/leu2Δ/ leu2Δ/leu2Δ/ura3Δ/ura3Δ/ ura3Δ/his3Δ/ his3Δ/ his3Δ/met15Δ/met15Δ/ met15Δ/TUP1::TAP/TUP1::TAP/TUP1::TAP* | | |
| YZ1606 | *MATa/a/a/a/leu2Δ/leu2Δ/leu2Δ/leu2Δ/ura3Δ/ura3Δ/ ura3Δ/ura3Δ/his3Δ/his3Δ/his3Δ/his3Δ/met15Δ/met15Δ/ met15Δ/met15Δ/NQM1::TAP/NQM1::TAP/NQM1::TAP/NQM1::TAP* | | |
| YZ1608 | *MATa/a/a/a/leu2Δ/leu2Δ/leu2Δ/leu2Δ/ura3Δ/ura3Δ/ ura3Δ/ura3Δ/his3Δ/his3Δ/his3Δ/his3Δ/met15Δ/met15Δ/ met15Δ/met15Δ/BRE2::TAP/BRE2::TAP/BRE2::TAP/BRE2::TAP* | | |
| YZ1618 | *MATa/a/a/a/leu2Δ/leu2Δ/leu2Δ/leu2Δ/ura3Δ/ura3Δ/ ura3Δ/ura3Δ/his3Δ/his3Δ/his3Δ/his3Δ/met15Δ/met15Δ/ met15Δ/met15Δ/RTG2::TAP/RTG2::TAP/RTG2::TAP/RTG2::TAP* | | |
| YZ1615 | *MATa/a/a/a/leu2Δ/leu2Δ/leu2Δ/leu2Δ/ura3Δ/ura3Δ/ ura3Δ/ura3Δ/his3Δ/his3Δ/his3Δ/his3Δ/met15Δ/met15Δ/ met15Δ/met15Δ/TUP1::TAP/TUP1::TAP/TUP1::TAP/TUP1::TAP* | | |
| | *MATa/a/a/a/leu2Δ/leu2Δ/leu2Δ/leu2Δ/ura3Δ/ura3Δ/ ura3Δ/ura3Δ/his3Δ/his3Δ/his3Δ/his3Δ/met15Δ/met15Δ/ met15Δ/met15Δ/ RFA1::TAP/RFA1::TAP/RFA1::TAP/RFA1::TAP* | | |
| YZ1847 | *BY4742 WT rho0: MATa his3Δ1 leu2Δ0 lys2Δ0 ura3Δ0* | | Gift from Prof. Dr. Herrmann |
| YZ1866 | *MATa his3Δ/ leu2Δ/ met15Δ/ ura3Δ/tup1::natNT2* | | |
| YZ1824 | *MATa his3Δ/ leu2Δ/ met15Δ/ ura3Δ/cln3:: natNT2* | | |
| YZ1859 | *MATa ura3Δ/ leu2Δ/ his3Δ/ met15Δ/ sch9::kanMX4* | | Euroscarf |
| YZ1860 | *MATa ura3Δ/ leu2Δ/ his3Δ/ met15Δ/ sch9::kanMX4/TUP1-6HA::natNT2* | | |
| YZ1862 | *MATa ura3Δ/ leu2Δ/ his3Δ/ met15Δ/ TUP1-6HA::natNT2* | | |
| Plasmid: | pRS414::sch9as (T492G) | A gift from Robbie Loewith | |

**Supplementary Table 2. List of used antibodies**

Type or paste caption here. Create a page break and paste in the Table above the

| Antigen | Name | Host species | Source | Note |
|---|---|---|---|---|
| Mrp10 | | Rabbit | | Gift from Prof. Dr. Herrmann |
| Mrpl40 | | Rabbit | | Gift from Prof. Dr. Herrmann |
| Oxa1 | | Rabbit | | Gift from Prof. Dr. Herrmann |
| Ilv5 | | Rabbit | | Gift from Prof. Dr. Herrmann |
| Mrps5 | | Rabbit | | Gift from Prof. Dr. Herrmann |
| Mdh1 | | Rabbit | | Gift from Prof. Dr. Herrmann |
| Clb2 | | Rabbit | | Gift from Dr. Zachariae |
| Rps23 | Sc-100837 | mouse | Santa Cruz | |
| Rpl9 | AP16409b-ev | Rabbit | ABGENT | |
| Sch9 | ab56203 | Rabbit | abcam | |
| PAP | P1291 | Rabbit | Sigma -Aldrich | |
| Anti-HA-Tag (F-7) | Sc-7392 | mouse | Santa Cruz | |
| Anti-Puromycin (clone 12D10) | MABE343 | mouse | Merck | |
| Rps21 | 16946-1-AP | Rabbit | Proteintech | |
| Rps8 | Ab201454 | Rabbit | abcam | |
| Rps19 | Sc-100836 | mouse | Santa Cruz | |
| Rpl22 | Sc-136413 | mouse | Santa Cruz | |
| Rpl5 | Ab86863 | Rabbit | abcam | |
| Rpl21 | Sc-393663 | mouse | Santa Cruz | |
| Tle1 | sc-137098 | mouse | Santa Cruz | |
| GAPDH (14C10) | 2118 | Rabbit | Cell signaling | |
| p70 S6 Kinase (49D7) | 2708 | Rabbit | Cell signaling | |
| Phospho-p70 S6 Kinase (Thr389) | 9205 | Rabbit | Cell signaling | |

**Supplementary Table 3 List of primers used for qPCR**

| Gene | Forward primer (sequence 5'-3') | Reverse primer (sequence 5'-3') |
|---|---|---|
| BRE2 | GTGTACGGGTATGGTATTAG | TATCACCCTCTTTCAATGGC |
| RTG2 | GCCTCTTATCCAAAGGAGTT | GCGCTCTAGCTCTGTGAGAT |
| TUP1 | CTATCTGACGATTCTGCTGC | GTGGTAGTAGTGGTTGTCATT |
| ILV5 | GGTTACGTTTACCAAACCAC | AGCCAAGAACATACCGTGG |
| MDH1 | CAACTCAGGAAAGGGTTAAC | TCTCTTGTCATCAGACATCAA |
| MRPL40 | GATTCCGTAGTGAGAAACCC | GCTATATCTTTCGCTGTCAAG |
| RPL9b | GAAACGTCCCAGTTAGAGAT | ATTTTGGGAAACGTCTTCAAC |
| IRS4 | TCTCTACATG AGGGAAATCTG | TGGTTCTGATCAGATTCAGAA |
| 5.8S | GTTGCGGCCATATCTACCAG | AGCACCTGAGTTTCGCGTAT |
| 25S | AGAGCCAATCCTTATCCCG | GGAAGCTCCGTTTCAAAGG |
| ACT1 | CACCCTGTTCTTTTGACTGA | CGTAGAAGGCTGGAACGTTG |
| FUS1 | TTCCAAACCCATAATACCGCA | CACTTTGGGGTCTAACGAAAT |
| HSP12 | AGGTAGAAAAGGATTCGGTGA | GGCCTTGTCAGTGATGTATTC |
| TMA10 | GACAGTCCACGAAGCAAAGT | GCCTTTCCCATAGCCTCCT |
| NCA3 | GATTTGGTGGATGGGCCTCT | GACATTCCAGGTTCACATGC |
| COX3 | TTGAAGCTGTACAACCTACC | CCTGCGATTAAGGCATGATG |
| 18S rRNA | GATGGTAGTCGCCGTGCC | GCCTGCTGCCTTCCTTGG |
| 28S rRNA | AGAGGTAAACGGGTGGGGTC | GGGGTCGGGAGGAACGG |

**Supplementary Table 4. Used software & packages for PloiDEx.**

| | | |
|---|---|---|
| MaxQuant v1.6.3.3 | Cox and Mann, 2008 | https://maxquant.net/ |
| Perseus v1.6.14 | Tyanova et al., 2016 | https://maxquant.net/perseus/ |
| R v3.6.1 | N/A | https://www.r-project.org/ |
| R package: ggplot2 v3.2.0 | Wickham H 2016 | https://ggplot2.tidyverse.org/ |
| R package: dplyr v0.8.3 | N/A | https://dplyr.tidyverse.org/ |
| R package: reshape2 v1.4.3 | N/A | https://cran.r-project.org/web/packages/reshape2/index.html |
| R package: ggalt | N/A | https://cran.r-project.org/web/packages/ggalt/index.html |
| R package: EQL v1.0-1 | N/A | https://cran.r-project.org/web/packages/EQL/index.html |
| R package: kdensity v1.0.1 | N/A | https://cran.r-project.org/web/packages/kdensity/index.html |
| YeastEnrichR | Chen et al., 2013 | https://maayanlab.cloud/YeastEnrichr |

**Supplementary Data 1. (separate file) Proteome data**

Excel table with proteome data set.

**Supplementary Data 2. (separate file) Transcriptome data**

Excel table with transcriptome data set.

**Supplementary Data 3. (separate file) Gene set enrichment analysis**

Excel table with results of the Gene set enrichment analysis.
